# Anti-activin treatment increases T cell infiltration in breast and pancreatic tumours and promotes survival in a SMAD4-null mouse pancreatic cancer model

**DOI:** 10.1101/2025.06.13.659133

**Authors:** Simon McCluney, Danielle Park, Daniel S.J. Miller, Merima Mehić, Robert D Bloxham, Emma De Vries, Xuelu Wang, Stefan Boeing, Luigi D’Antonio, Łukasz Wieteska, Gwen Pyeatt, Stephanie Strohbuecker, Probir Chakravarty, Taiana Maia De Oliveira, Robert W Wilkinson, Marko Hyvönen, James Hunt, Caroline S Hill

## Abstract

Activin A and B share the same downstream signalling pathway (activation of SMAD2/3) as TGF-β and consequently elicit many of the same functional responses as TGF-β, including immune suppression, activation of cancer-associated fibroblasts (CAFs) and extracellular matrix production and remodelling. However, activin’s role in tumourigenesis has been relatively overlooked compared to TGF-β’s. We generated and characterized a dual specificity human antibody that recognizes both activin A and B and compared its activity in syngeneic mouse models of breast cancer and pancreatic ductal adenocarcinoma (PDAC) with an activin A-specific antibody. We demonstrate that activin A and B are central to the function of CAFs and therapeutic inhibition of activin results in a reduction of collagen rich desmoplastic barriers, enabling the infiltration of cytotoxic T cells. This is correlated with an upregulation of the T cell chemoattractant CXCL10, which is normally repressed by activin signalling. Interestingly, despite greater T cell infiltration, activin A inhibition resulted in poorer survival in the KPC mouse model of PDAC and slightly larger tumours in the breast cancer model, indicating a tumour suppressive role of activin A-rich CAFs. Strikingly, however, treatment with the same anti-activin A antibody of PDAC tumours where SMAD4 is deleted in the tumour cells, resulted in increased survival, which was potentiated with additional treatment with immune checkpoint inhibitors. These results suggest that anti-activin therapy has potential for the cohort of PDAC patients exhibiting inactivation of SMAD4.

## Introduction

The transforming growth factor β (TGF-β) ligands (TGF-β1, 2 and 3) have long been known to play important roles as both tumour suppressors and tumour promoters in many different types of human cancer. As a tumour suppressor, TGF-β can induce apoptosis, particularly in tumours containing KRAS mutations ^1,2^, induce cytostatic effects ^3^, and also can elicit indirect effects via the tumour stroma, primarily by preventing inflammation ^4^. As tumours develop, cancer cells use various strategies to circumvent the tumour suppressive effects of TGF-β, such as by mutating or deleting key components of the TGF-β pathway to make the cancer cells unresponsive to ligand. Loss-of-function mutations are found in the type I or type II receptors (TGFBR1 and TGFBR2 respectively), or in the SMADs which transduce the signal to the nucleus. In particular, SMAD4, the so-called common mediator SMAD, which is required for most TGF-β responses ^5,6^, is deleted or acquires loss-of-function mutations in about a third of pancreatic cancers as well as in a subset of colorectal cancers ^7–11^. As a tumour promoter, TGF-β has numerous effects on the tumour stroma by reprogramming fibroblasts to cancer-associated fibroblasts (CAFs), which become a source of further TGF-β and activin (another subgroup of the TGF-β family) ligands, as well as of inflammatory cytokines ^6,12^. Moreover, TGF-β can act as a pro-angiogenic factor, and as an immune suppressor, by suppressing proliferation and cytolytic functions of cytotoxic T cells and preventing their access to the core of the tumour ^5^. TGF-β can also promote pro-tumourigenic phenotypes of tumour-associated macrophages and neutrophils ^5^.

Given these well documented tumour promoting activities, there have been numerous attempts to target TGF-β signalling therapeutically in cancer using, for example, ligand traps, neutralizing antibodies, or small molecule ATP competitive inhibitors of the type I receptor kinase activity ^3,6,13,14^. However, despite more than 20 molecules being tested in clinical trials over the past two decades, no treatment in solid tumours has yet been approved. More recently, there has been growing interest in combining anti-TGF-β therapy with immune checkpoint inhibitors (ICIs) such as anti-PD-L1 antibodies, in view of TGF-β’s strong immune-suppressive effects and the link of TGF-β signalling to poor response to ICIs ^15^. In one case this has been attempted with a bi-functional agent, M7824, comprising a TGF-β ligand trap with an anti-PD-L1 antibody ^3,6,13^. Although these combinations were very successful in preclinical mouse models ^16–18^, this approach has not yet translated to success in the clinic ^3,6,19^.

One issue with targeting the TGF-β ligands in disease, is that they share a downstream signalling pathway with other TGF-β family ligands, in particular, activin A and activin B (encoded by the genes *INHBA* and *INHBB* respectively), which are also widely expressed in many different tumours ^20^. Although the activin receptors (ACVR1B/ACVR1C and ACVR2A/ACVR2B) are distinct from the TGF-β receptors (TGFBR1 and TGFBR2), their activation by ligand results in phosphorylation of the same receptor-regulated SMADs, SMAD2 and SMAD3. These phosphorylated SMADs form complexes with SMAD4, which accumulate in the nucleus to transcriptionally regulate target genes. As a result, activin and TGF-β share many of the same target genes ^21^ and thus the presence of phosphorylated SMAD2/3 or a specific target gene signature cannot be used in vivo as an indication of the activity of a specific ligand ^3^. Furthermore, these ligands exhibit distinct signalling dynamics. Cell responses to the TGF-β ligands quickly attenuate due to rapid internalisation of receptor complexes, resulting in depletion of receptors from the cell surface ^22^. This means that long-term stimulation with TGF-β results in low levels of phosphorylated SMAD2/3. The activin signal, in contrast, is integrated over time. Thus, long-term activin stimulation results in high levels of phosphorylated SMAD2/3 ^23^. As a result, some activities that have been ascribed *in vivo* to the TGF-βs may be the result of activin signalling.

This view is supported by the fact that activities, such as immune suppression and CAF activation are co-induced by TGF-β and activin A and B ^20,24,25^. Moreover, loss-of-function mutations/deletions of activin receptors (ACVR1B, ACVR2A and to a lesser extent ACVR2B) are observed in pancreatic ductal adenocarcinoma (PDAC) as an alternative to SMAD4 mutations/deletions ^25–27^. This strongly suggests that activins have a tumour suppressor function in human PDAC. In addition, a number of pre-clinical studies in mice have shown that activins also drive tumorigenesis in PDAC, breast cancer, melanoma and other skin cancers ^28–36^ and is important for lung metastasis in breast cancer patients ^37^. Therefore, like TGF-β, evidence is emerging for activins playing both tumour suppressive and tumour promoting roles. Untangling these opposing roles is going to be essential for treatment of human cancer.

To delineate the roles specifically played by activins in tumourigenesis we developed and characterized a human dual specificity anti-activin A/B antibody of high affinity and compared it in syngeneic mouse models with an anti-activin A-specific antibody containing the variable regions of garetosmab ^38^ and with an IgG control. We show that these two anti-activin antibodies bind at different sites on mature activin A, but both effectively neutralize activin signalling in vivo. We chose to focus on two cancer types – invasive breast cancer which is associated with high levels of circulating activin A in patients with primary tumours and bone metastases ^39–41^, and PDAC, one of the hardest to treat cancers which is known to exhibit loss-of-function mutations in activin receptors or SMAD4, and where high levels of activin expression are associated with a worse prognosis ^25,27,32,42^. Using these antibodies, we demonstrate that the major effect of inhibiting activin A/B signalling in breast cancer is to allow infiltration of cytotoxic T cells into the core of the tumour and to decrease collagen deposition by CAFs. In the PDAC model, we show that neutralizing activins in tumours where SMAD4 is wild type results in poorer survival. Strikingly however, in mice bearing tumours in which SMAD4 has been deleted specifically in the tumour cells, inhibiting activin A function promotes survival, and this is further increased with treatment with anti-PD-L1 antibodies. These data indicate that we can successfully disentangle the tumour promoting and tumour suppressive effects of activin and suggest a therapeutic potential of inhibiting activin A signalling in human PDAC tumours that have lost SMAD4 function.

## Results

### *INHBA* and *INHBB* are strongly expressed in breast and pancreatic cancer and high expression of *INHBA* correlates with poorer survival

To confirm the relevance of activin expression in human cancer we analysed cancer genome atlas (TCGA) data. Patients with PDAC tumours expressing high levels of *INHBA* (which encodes activin A) showed reduced survival compared to those expressing low levels (Figure 1A). We followed this up by analysing levels of activin A and activin B protein in serum from patients with PDAC compared with age- and sex-matched normal controls. Both activin A and activin B levels were significantly enhanced in the serum from cancer patients compared with the controls (Figure 1B). To determine which cells in PDAC tumours expressed *INHBA* and *INHBB* (which encodes activin B), we reanalysed human single cell RNA-sequencing (scRNA-seq) data from PDAC tumours and control pancreases from patients with non-pancreatic tumours or non-malignant pancreatic tumours ^43^. Little expression of *INHBA* was observed in the control pancreases, but high expression was detected in CAFs, predominantly in the myCAF population ^12,44^ and in myeloid populations, particularly in macrophages (Figure 1C–E). In contrast, *INHBB* was expressed at low level in a range of cell types in both the control pancreases and in the PDAC tumours (Figure 1C). These findings were confirmed by performing combined RNAscope™ in situ hybridization and immunofluorescence for α smooth muscle actin (αSMA, encoded by the *ACTA2* gene), a marker of myCAFs ^12^. Much of the expression of *INHBA* co-localized with αSMA in human PDAC samples, whilst expression of *INHBB* was more uniform throughout the sample in epithelial cells and in CAFs (Figure 1F).

**Figure 1.**
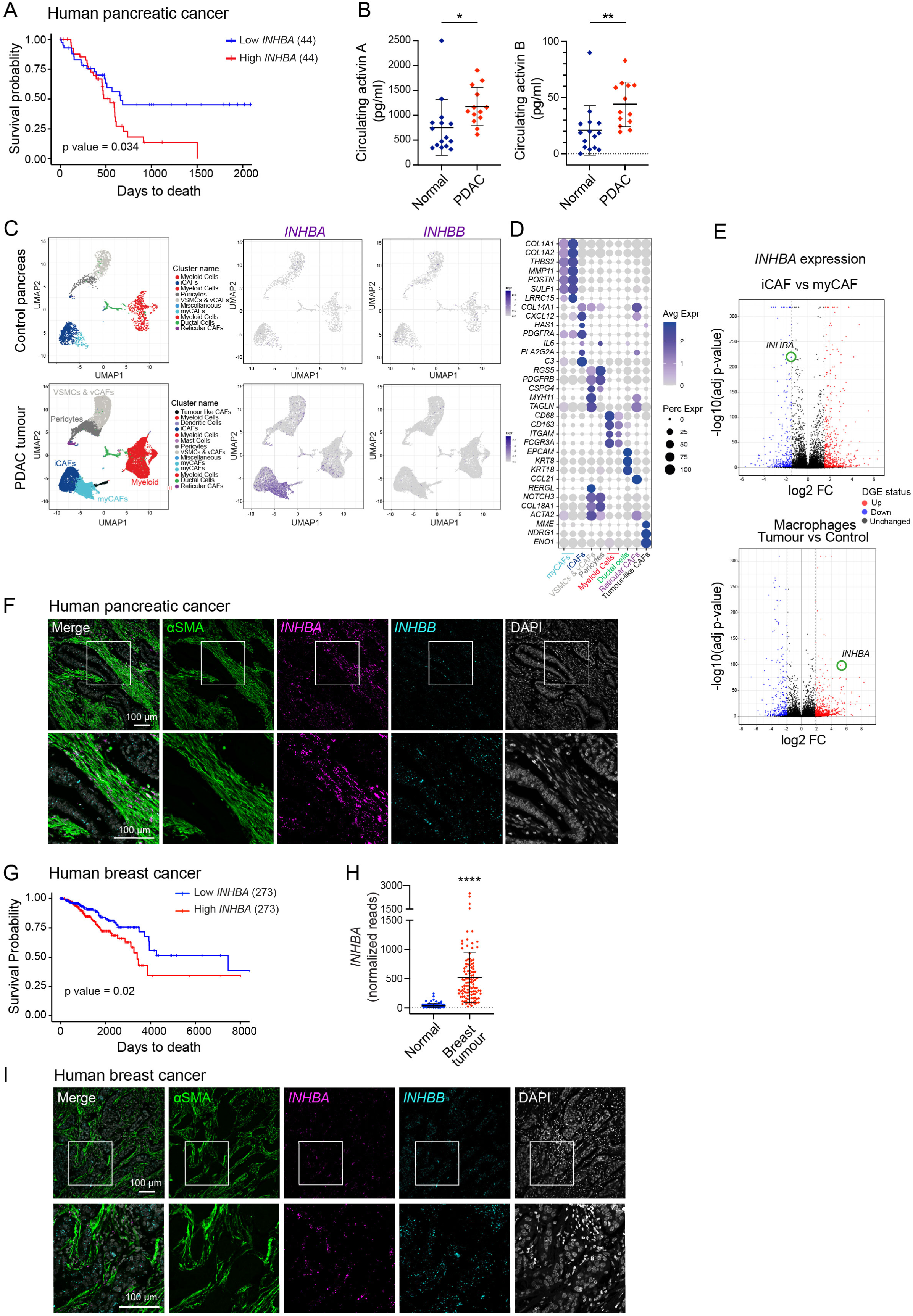
Activin A and activin B are expressed in human pancreatic and breast tumours, with activin A expression correlating with poor survival. (**A**) Kaplan Meier survival plots for PDAC patients (data from The Cancer Genome Atlas (TCGA)) stratified according to expression of *INHBA*. High and low expression defined as upper and lower quartiles respectively (44 tumours in each) from 176 tumours. The significance of the difference in survival is indicated by p = 0.034. (**B**) Circulating activin A and B expression in human PDAC patients and age and sex-matched healthy volunteers analysed by ELISA. Means ± SD are shown. *, p <0.05; **, p <0.01. (**C**) UMAP projections from a re-analysed scRNA-seq dataset from human pancreatic cancer and control pancreases from patients with non-pancreatic tumours or non-malignant pancreatic tumours ^43^. Subclustering of the original dataset to display fibroblast and myeloid populations. *INHBA* and *INHBB* expression is shown within subclustered populations for both control pancreases and PDAC samples. (**D**) Dot plot demonstrating representative marker genes across cell types. Dot size is proportional to the fraction of cells expressing specific genes. Colour intensity corresponds to the relative expression of specific genes. (**E**) Volcano plot demonstrating differential *INHBA* expression in iCAF vs myCAF populations (upper panel) and volcano plot demonstrating differential *INHBA* expression in tumour vs control samples within the myeloid cell cluster (lower panel). (**F**) Confocal microscopy images of human PDAC samples stained for *INHBA* mRNA (magenta) and *INHBB* mRNA (cyan) by RNAscope™ with immunostaining for αSMA (green) and DAPI staining to visualize the nuclei (grey). The bottom row shows higher magnifications of the regions marked with a white box. Scale bars correspond to 100 µm. (**G**) Kaplan Meier survival plots for breast cancer patients (data from The Cancer Genome Atlas (TCGA)) stratified according to expression of *INHBA*. High and low expression defined as upper and lower quartiles respectively (273 tumours in each) from 1092 tumours. The significance of the difference in survival is indicated by p = 0.02. (**H**) Graphs demonstrating *INHBA* transcript reads in human breast cancer tumour samples and healthy controls (data from TCGA). ****, p <0.001. (**I**) Confocal microscopy images of human breast cancer samples stained for *INHBA* mRNA (magenta) and *INHBB* mRNA (cyan) by RNAscope with immunostaining for αSMA (green) and DAPI staining to visualize the nuclei (grey). The bottom row shows higher magnifications of the regions marked with a white box. Scale bars correspond to 100 µm.

We next investigated whether a mouse model of PDAC (the KPC model ^45^) recapitulated our findings in human PDAC and found this to be the case. Reanalysis of scRNA-seq data from this mouse model demonstrated high levels of *Inhba* expression in myeloid cells, myCAFs and low expression in ductal cells, whilst *Inhbb* transcripts were detected in myCAFs, iCAFs and ductal cells but were absent in cells of the myeloid lineage (Figure S1A–C). RNAscope™ for *Inhba* or *Inhbb* with immunofluorescence for αSMA confirmed the scRNA-seq analysis with *Inhba* transcripts co-localising with αSMA staining, whilst expression of *Inhbb* was more delocalized (Figure S1D).

We went on to investigate whether other tumour types also expressed high levels of these two activins, focusing on human invasive breast cancer. Analysis of TCGA data indicated that high *INHBA* expression was associated with poorer survival and *INHBA* expression was significantly higher in tumours compared with normal mammary tissue (Figure 1G, H). RNAscope™ for *INHBA* or *INHBB* with immunofluorescence for αSMA indicated that, like PDAC, expression of *INHBA* was mainly associated with CAFs, whilst *INHBB* was more widely expressed (Figure 1I). A similar analysis in a syngeneic mouse orthotopic transplantation model of breast cancer (D2A1-m2 ^46^) indicated very high levels of *Inhba* expression within CAFs and in tumour cells, but much lower levels of *Inhbb* expression (Figure S1E). Analysis of mouse plasma indicated increased levels of activin A and activin B in tumour-bearing mice compared with wildtype mice in the MMTV-PyMT model ^47^ and in the D2A1-m2 model (Figure S1F, G).

In summary, both human and mouse PDAC and breast tumours exhibit high levels of *INHBA* expression predominantly in the myCAFs and myeloid populations, as well as broad expression of *INHBB*. In both cancer types, raised levels of circulating activin A and B in the bloodstream are detected in tumours versus controls.

### Activin secreted by CAFs promotes ECM production and contractility

Building on our observation that CAFs are a major source of activins, we investigated their functional role in this cell type. We first determined whether activins were the only TGF-β family member secreted by the CAFs. Conditioned media was prepared from an immortalized clone of CAFs (CAF1), obtained from the MMTV-PyMT mouse breast cancer model ^48^ and from immortalized normal fibroblasts (NF) extracted from normal mammary glands. This was used to induce phosphorylated SMAD2 (pSMAD2) in HaCaT cells (Figure 2A, B). The CAFs, but not the NFs, secreted an activity that could be inhibited by the activin A and B antagonist, follistatin, but not a TGF-β blocking antibody, indicating that the CAFs secrete activin, but not TGF-β (Figure 2B).

**Figure 2.**
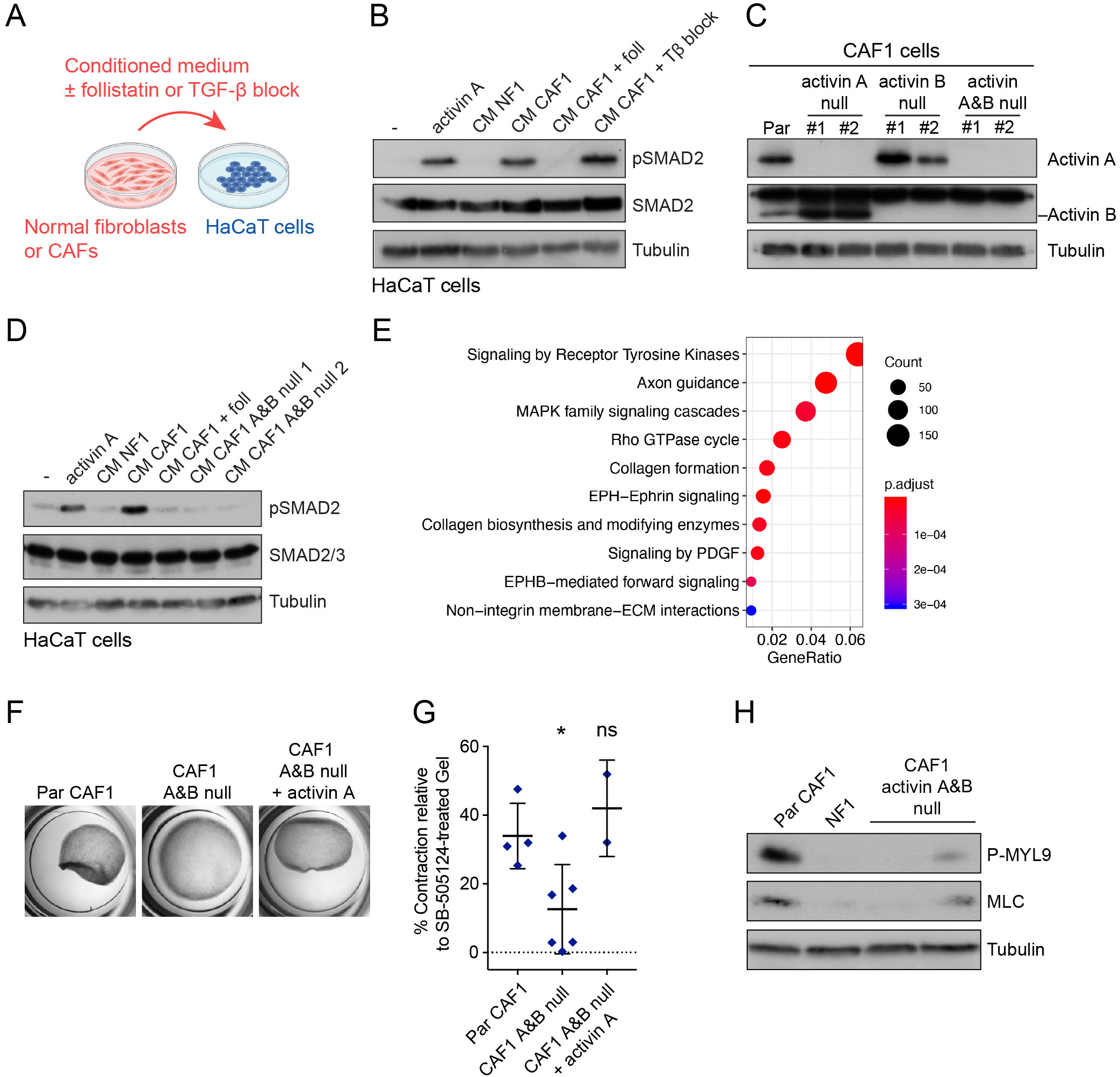
CAFs produce activin A and activin B which are required for collagen gel contraction and extracellular matrix production. (**A**) Schematic illustrating the process of using conditioned media from normal fibroblasts or CAFs to stimulate naïve HaCaT cells with various antagonists to study TGF-β family pathway activation. (**B**) Western blot demonstrating induction of TGF-β family signalling via pSMAD2 activation. Recombinant activin A ligand (20 ng/ml) is used as positive control. HaCaT cells were stimulated for 1 hr with conditioned media (CM) from normal fibroblasts (NF1) or from CAFs with or without 500 µg/ml follistatin (foll) to inhibit activin signalling or 1 µg/ml TGF-β blocking antibody (Tβ block). Whole cell extracts were blotted for pSMAD2 with SMAD2, and Tubulin used as loading controls. (**C**) Western blot characterizing the CAF1 parental line (Par) and CAF1 cells knocked out for activin A, activin B knockout lines or both. Extracts from 2 clones of the knockout cells were analysed (#1 and #2) and were blotted with the antibodies shown. Tubulin was used as a loading control. (**D**) As for (B) but conditioned media was prepared from NF1, CAF1 cells and from CAF1 cells deleted for both activin A and B. (**E**) Bulk RNA-seq was performed where the transcripts from the parental (Par) CAF1 were compared with the CAF1 cells deleted for activin A and B. Significantly differentially expressed genes between the two cell types were analyzed by Reactome pathway analysis and the hits were ordered according to GeneRatio with the highest at the top of the list. (**F, G**) Collagen contraction assay of parental CAF1 cells (Par), the CAF1 cells deleted for both activin A and B with or without addition of recombinant activin A (20 ng/ml). Cells were embedded in collagen gel and their ability to contract was measured on day 3 (F). Quantification is shown (G) where the % contraction of respective gels at day 3 is plotted relative to samples treated with 10 µM SB-505124 where activin signalling has been abolished. Means ± SD are shown. The p values are shown relative to the parental cells. ns, not significant; *, p < 0.05. (**H**) Western blot demonstrating levels of P-MYL9 and total MLC in parental CAF1 cells, NF1 cells and 2 separate clones of activin A and B-null CAF1 cells. Tubulin is used as a loading control.

To investigate the functional role of activins in the CAFs, we generated single and double knockouts of activin A and activin B using CRISPR/Cas9 (Figure 2C) and confirmed that the double knockouts did not secrete any ligands that could induce pSMAD2 in HaCaT cells (Figure 2D). Reactome pathway analysis of bulk RNA-seq data, comparing parental CAF1 cells with those deleted for activin A and B, revealed collagen biosynthesis and collagen formation as major pathways, among others, that differed between the two cell types (Figure 2E). We therefore investigated whether activin signalling was required for the CAF’s ability to remodel the extracellular matrix using the collagen gel contraction assay. Parental CAF1 cells could readily contract collagen, but the activin A and B-null CAF1 cells were unable to, and this deficit could be rescued by addition of recombinant activin A (Figure 2F, G). Consistent with this, the activin A and B-null CAF1 cells expressed considerably lower levels of phosphorylated MYL9, which is required for contraction, compared to the parental line (Figure 2H). Thus, activin A and B produced by CAFs is required for ECM production and CAF contractility in vitro.

### Generation and characterization of anti-activin neutralizing antibodies

To determine the role of the secreted activin A and B in vivo we generated a dual specificity anti-activin A/B antibody by screening AstraZeneca’s phage display libraries of randomly combined human single chain variable fragments (scFvs) using recombinant mature activin A and activin B as baits. This resulted in a dual specificity anti-activin A/B antibody that was converted to the human IgG1 format. Using bio-layer interferometry we demonstrated that this antibody has a K_D_ for human activin A of 130 pM and a K_D_ for human activin B of 750 pM (Figure 3A, B). We demonstrated that the anti-activin A/B antibody could specifically inhibit the activity of both human activin A and activin B in a CAGA_12_-luciferase assay in HEK293T cells (Figure 3C, D). Consistent with high conservation between human and mouse activins (100% and 97% identify for activin A and B, respectively), the antibody could also effectively block the activity of the mouse ligands as it inhibited the ability of activin A and B secreted by mouse CAFs to activate the phosphorylation of SMAD2 in HaCaT cells (Figure 3E). As a comparator for our in vivo assays, we also produced a high affinity anti-activin A antibody in the same IgG1 format as our anti-activin A/B antibody, using the variable regions of garetosmab ^38^. This antibody was specific for activin A only and had a higher affinity for activin A (∼8.7 pM, measured by surface plasmon resonance) compared with the anti-Activin A/B antibody (Figure 3C, D; S2A, D and G). In addition, we checked the specificity of these anti-activin antibodies against other TGF-β family ligands. Neither antibody exhibited any affinity for TGF-β3 (Figure S2C, F, I). The dual specificity anti-activin A/B antibody showed a very weak affinity for BMP2 (K_D_ = 0.102 µM) which was approximately 500 times weaker than its affinity for activin A in the same assay (K_D_ = 0.254 nM) (Figure S2B, H), and much weaker than the affinity of BMP2 for its cognate type I receptors ^49^. The garetosmab-like anti-activin A antibody showed no detectable binding to BMP2 (Figure S2E).

**Figure 3.**
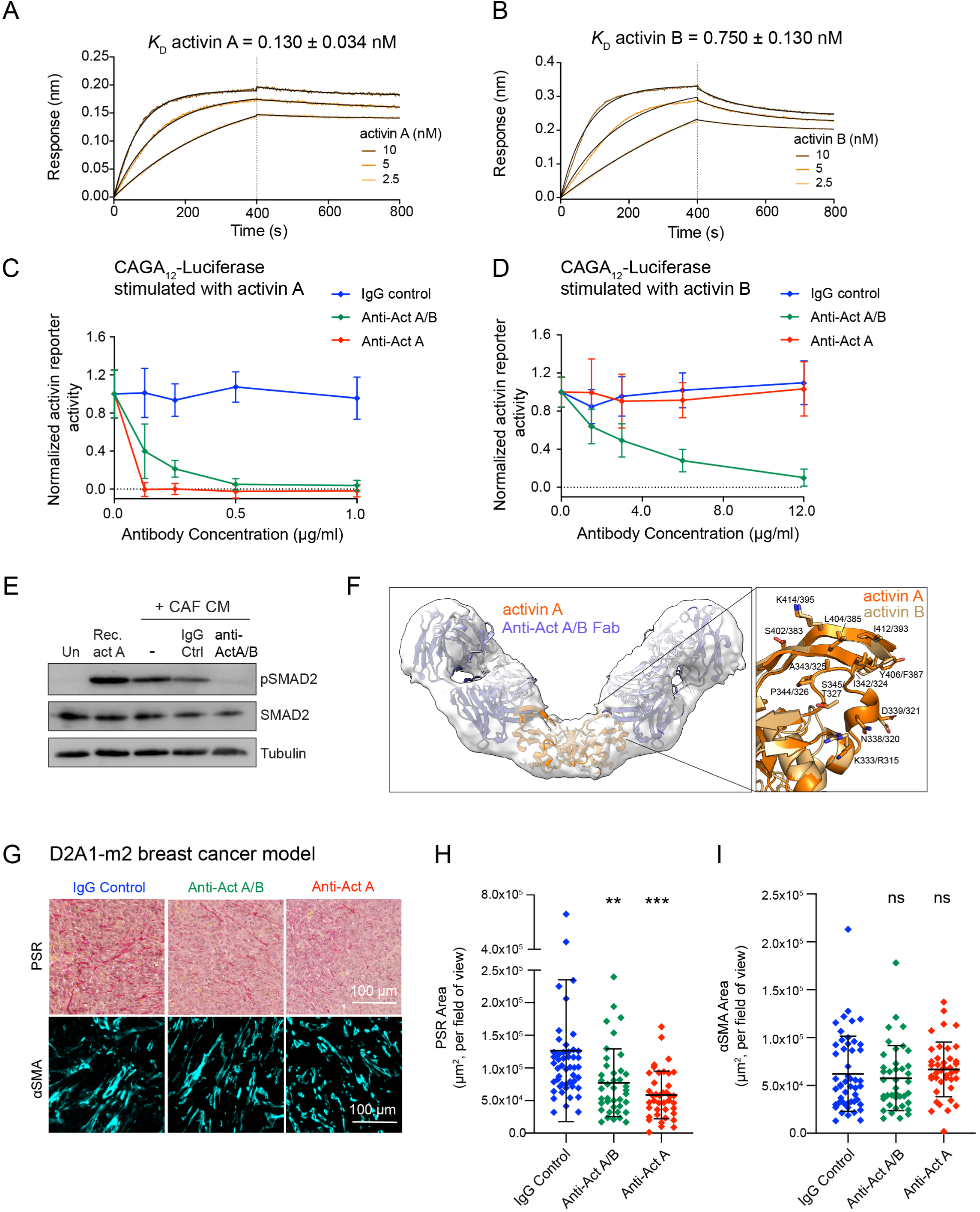
Generation and characterization anti-activin antibodies. (**A, B**) The binding affinity of anti-activin A/B antibody to activin A (A) and activin B (B) was measured by bio-layer interferometry. (**C, D**) Specificity of the anti-activin A/B antibody compared with an anti-activin A antibody and IgG control antibodies. HEK293T cells stably containing the activin reporter CAGA_12_-Luciferase and an internal TK-Renilla control were stimulated with either activin A ligand (C) or activin B ligand (D) in the presence of different concentrations of the antibodies as indicated. Normalized luciferase values are plotted relative to the Renilla control. Means and ± SD shown. (**E**) Western blot demonstrating that the anti-activin A/B antibody inhibits induction of pSMAD2 by conditioned media (CM) from mouse cancer associated fibroblasts (CAFs) which is shown to contain activin A and activin B, but not TGF-β (see Figure 2B). HaCaT cells were stimulated for 1 hr with CM from CAFs with either a control antibody or anti-Act A/B (200 µg/ml), or no addition. Recombinant activin A (Rec act A) (20 ng/ml) ligand was used as positive control. Whole cell extracts were blotted for pSMAD2 with SMAD2, and Tubulin used as loading controls. (**F**) Left panel, structure of activin A:anti-activin A/B Fab complex. AlphaFold 3 model of mature activin A (orange) with anti-activin A/B Fab fragment (light and dark blue) fitted into the cryo-EM map of the complex at 5.4 Å. Right panel, interaction site of anti-activin A/B on activin A and B. AlphaFold 3 models of mature activin A (bright orange) and B (lighter orange) superimposed on one protomer of the dimeric ligand. Residues that have side chain atoms within 4.0 Å from the anti-activin A/B Fab fragment (which are not shown in the figure) are shown as sticks and labelled. Numbering of the residues are as per the Uniprot entries of the two proteins. (**G**) Representative images of tumours from BALB/c mice injected with D2A1-m2 cells and treated with IgG control, anti-activin A/B or anti-activin A antibodies stained with picrosirius red (PSR - red staining) for collagen deposition and for αSMA to label fibroblasts. Scale bar corresponds to 100 µm. (**H**) Quantification of picrosirius red staining in tumours from BALB/c mice injected with D2A1-m2 cells and treated with IgG control, anti-activin A/B or anti-activin A antibodies. Data are represented as % of total positive cells, per field of view. 10 fields of view are shown from 5 tumours (IgG) and 4 tumours (anti-activin A/B and anti-activin A samples). Means ± SD are shown. ****, p <0.0001. (**I**) Quantification of αSMA staining in tumours from BALB/c mice injected with D2A1-m2 cells and treated with IgG control, anti-activin A/B or anti-activin A antibodies. Data are represented as % of total positive cells, per field of view. The same tumours were analysed as in H. Means ± SD are shown. ns, not significant. In H and I, the statistical significance is calculated relative to the IgG control.

To determine how the anti-activin A/B antibody bound to activin A we generated a complex of the Fab fragments of this antibody with mature activin A for cryo-EM analysis. Structural characterization of the complex was hampered by the structural heterogeneity of the activin A dimer ^50^. We were therefore able only to generate a map at 5.4 Å resolution, which nevertheless shows a clear V-shaped structure, consistent with Fab fragments binding to the so-called knuckle epitopes where the activin type II receptors bind. To gain further insight, we generated structural models of anti-activin A/B Fab fragments in complex with mature activin A and activin B using the AlphaFold3 server (https://alphafoldserver.com). All models generated for each complex by AlphaFold3 were fully consistent with each other (data not shown), as were the models using either activin A or activin B as the ligand. The activin A:anti-activin A/B Fab models showed a perfect fit to the cryo-EM map (Figure 3F). The interfaces between the anti-activin A/B antibody and activin A or B in the AlphaFold3 models were largely overlapping with the known binding site of the ACVR2B extracellular domain (Figure S2J). We validated these findings with bio-layer interferometry that showed that the extracellular domain of the activin type II receptor, ACVR2B, as well as the activin antagonist follistatin ^21,51,52^ competed with the anti-activin A/B antibody for activin A binding (Figure S2L). Analysis of the residues predicted in activin A and B to contact the anti-activin A/B antibody showed almost perfect conservation, consistent with its ability to inhibit both growth factors (Figure 3F; S2K). These binding site residues are more poorly conserved between the activins and BMP2 or TGF-β3 (Figure S2K), in agreement with our specificity data. Interestingly, sequential binding data (Figure S2M) suggests non-overlapping epitopes for the two antibodies, which is in agreement with the high confidence AlphaFold3 models for their complexes with activin A (Figure S2N).

### Validating the anti-activin antibodies in vivo

Having fully characterized the antibodies in vitro, we validated them as neutralizing antibodies in vivo. The stability of the two anti-activin antibodies and the IgG control in the mouse circulation was similar, with a drop of about 50% measured after 7 days (Figure S3A). We then investigated their pharmacodynamics, measuring their ability to interact with their targets and elicit a biological effect. To assess the binding of the antibodies to circulating activin, we performed ELISAs for activin A in the blood of mice injected with an IgG1 control or the anti-activin antibodies. We detected a significant increase in the amount of activin A in the bloodstream of anti-activin-treated mice, compared with the IgG control (Figure S3B). We interpreted this as resulting from activin–antibody complexes forming in the blood which are subsequently disrupted in the ELISA assay performed in denaturing conditions. This phenomenon has been seen in similar assays from normal human subjects treated with anti-activin therapy ^38^. To determine whether the anti-activin antibodies could neutralize activin activity in tumours, we stained D2A1-m2 breast tumours from mice treated with IgG control or the anti-activin antibodies for pSMAD2, the downstream effector of the activin pathway ^53^. We observed a significant decrease in the amount of pSMAD2 in tumours treated with anti-activin A/B or anti-Activin A compared with IgG control, indicating that the antibodies are penetrating the tumours and blocking activin activity (Figure S3C, D).

### Anti-activin treatment of mice results in decreased collagen production and increased infiltration of cytotoxic T cells into breast tumours

Having demonstrated that both anti-activin antibodies had effective neutralizing activity in vivo, we investigated the effect of inhibiting activin A and B *in vivo* on tumour development, beginning with a metastatic breast cancer model where D2A1-m2 cells were orthotopically injected into syngeneic BALB/c mice. The resulting tumours are rich in CAFs, express high levels of *Inhba* and are immune excluded (ref ^54^; Figure S1E). We first investigated the effect of anti-activin treatment on collagen levels in tumours, in view of our in vitro data showing that activin A and B are required for CAF contractility and ECM production. We showed that treatment of mice with either of the anti-activin antibodies depleted collagen levels in the tumours as measured by picrosirius red staining compared with the IgG control (Figure 3G, H). However, we saw no effect on the density of CAFs as measured by αSMA staining (Figure 3I).

To gain a more detailed understanding of how anti-activin therapy affects the tumour microenvironment (TME), we performed scRNA-seq on tumours from D2A1-m2 injected mice treated with anti-activin A/B, anti-activin A or IgG control antibodies (Figure 4A–C). In the D2a1-m2 model the T cells are typically restricted to the stromal rich boundary at the tumour periphery ^54^. We observed a substantial increase in the proportion of infiltrating T cells (both CD4^+^ and CD8^+^) in the tumours of mice treated with anti-activin antibodies, compared with those treated with the IgG control, which agreed with an inverse correlation between a CD8^+^ T cell signature and *INHBA* expression in human breast cancer TCGA datasets (Figure S4A). To determine whether anti-activin treatment was facilitating the conversion from immune excluded to immune ‘hot’ or infiltrated we performed immunohistochemistry to assess the spatial resolution of T cells. We observed a significant increase in CD4^+^ and CD8^+^ T cells in mice treated with anti-activin therapy, particularly within the tumour core (Figure 4D, E). These data suggest that anti-activin treatment facilitates the recruitment of T cells into the tumour, switching them from an immune excluded to a more immune hot phenotype. We also investigated B cells as the scRNA-seq analysis suggested that anti-activin A/B treatment resulted in increased infiltration of B cells. However, we found no significant increase in B cell infiltration with anti-activin therapy (Figure 4F). The scRNA-seq data also revealed which cells in the tumour expressed *Inhba* and *Inhbb*. Very little expression of *Inhbb* was detected, consistent with our RNAscope™ data, but *Inhba* was strongly expressed in cancer cells, macrophages and neutrophils and other undefined stromal cells (Figure S4B; S1E). Given that fibroblasts were not efficiently extracted after tumour dissociation we were unable to use the scRNA-seq dataset to visualize the *Inhba* or *Inhbb* expression in CAFs.

**Figure 4.**
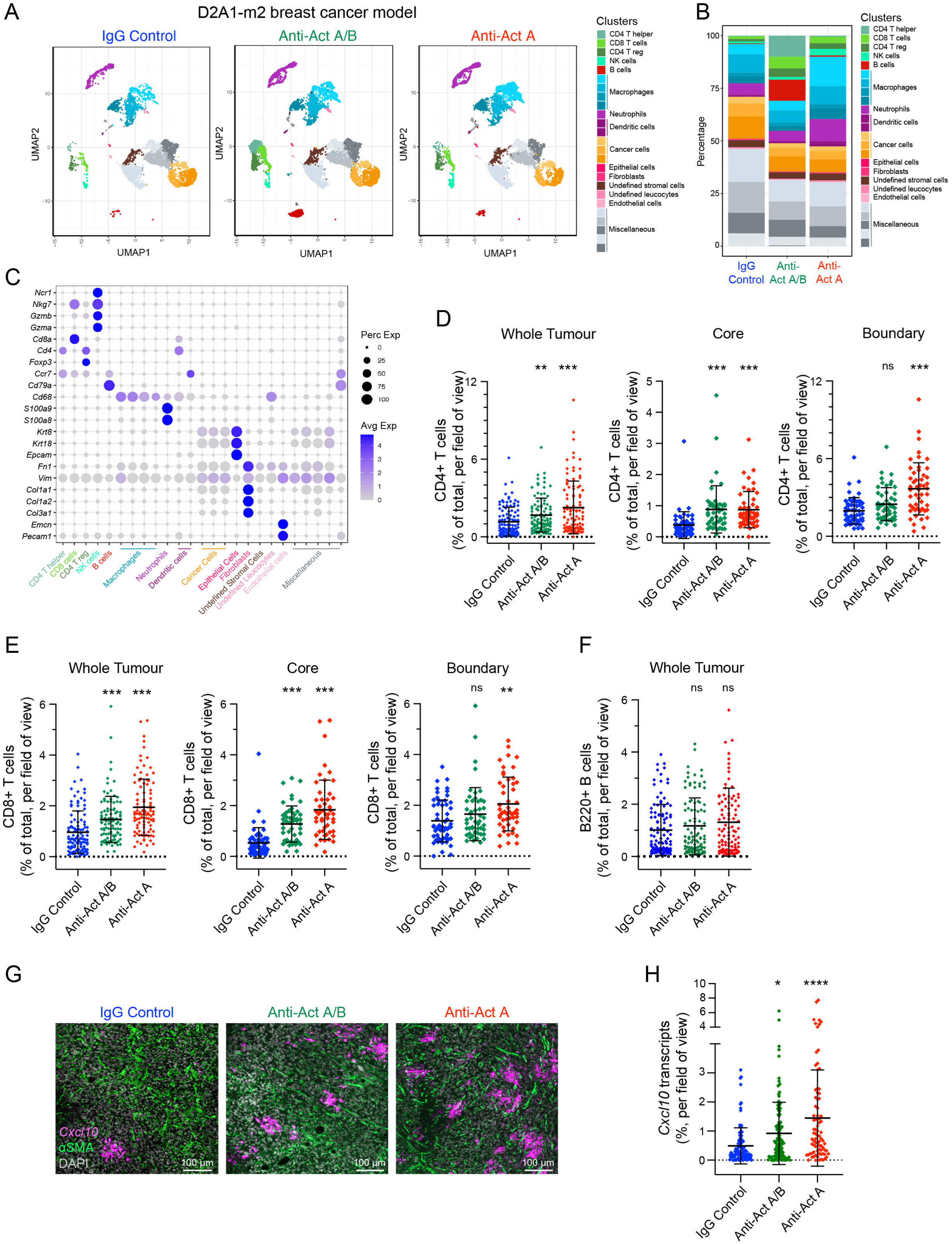
Treatment of breast tumours with anti-activin antibodies enhances infiltration of CD4^+^ and CD8^+^ T cells. (**A**) UMAP projections from an scRNA-seq dataset of tumours from the D2A1-m2 mouse breast cancer model (D2A1-m2 injected into the mammary fat pad of BALB/c mice). Mice were treated with IgG control, anti-activin A/B or anti-activin A antibodies. Cells are coloured according to cell type. (**B**) Area plots demonstrating percentage of cells per cell type across the three treatment groups; IgG control, anti-activin A/B or anti-activin A antibodies. (**C**) Dot plot demonstrating representative marker genes across cell types. Dot size is proportional to the fraction of cells expressing specific genes. Colour intensity corresponds to the relative expression of specific genes. (**D–F**) Quantifications of immune cell infiltration in tumours from the mouse breast cancer model (D2A1-m2) where mice were treated with IgG control, anti-activin A/B or anti-activin A antibodies. CD4^+^ (D), CD8^+^ (E) and B220^+^ (F) cells were detected by immunostaining and quantified in tumours as percentage of cells of interest per field of view. 10 fields of view were quantified per tumour. For the CD4+ and CD8+ cells, the whole tumour was quantified (n=11 for the IgG condition and n=10 for the other two conditions), and the quantification was additionally subcategorised with core and boundary quantification. For B220^+^ cells the whole tumour was quantified (n=6 for each condition). ns, not significant; **, p <0.01; ***, p <0.001. (**G**) Representative confocal microscopy images demonstrating staining of mouse tumours of the D2A1-m2 model treated with IgG control, anti-activin A/B or anti-activin A antibodies for *Cxcl10* mRNA by RNA scope (magenta) αSMA protein by immunofluorescence (green) and DAPI to mark the nuclei (grey). Scale bar represents 100 µm. (**H**) Quantifications of *Cxcl10* mRNA staining in the experiment shown in (G). Data are represented as % of positive area, per field of view. *, p <0.05; ****, <0.0001. 40 fields of view per tumour. IgG control (n=3), anti-activin A/B (n=3), anti-activin A (n=2). In all cases, the statistical significance is calculated relative to the IgG control.

We reasoned that the anti-activin therapy might increase cytotoxic T cell infiltration partly by reducing the collagen rich desmoplasia that physically excludes immune cells (Figure 3G, H), but also as a result of increased expression of cytotoxic T cell chemoattractants ^55^. We investigated this further by mining the RNA-seq data from the CAF analysis to determine what chemokines were upregulated in CAFs when activin A and B were deleted (Figure S4C). There were several candidates, which we validated by qPCR (data not shown), and found that the most reproducibly upregulated chemokine in activin A and B-null CAFs compared with the parental CAFs was *CXCL10* (Figure S4D). CXCL10 has been shown previously to be a key chemokine expressed in the TME that promotes CD8^+^ T cell infiltration by signalling through its receptor CXCR3 on activated CD8^+^ T cells ^56,57^. The enhanced expression of *CXCL10* was validated using RNAScope™ in tumours from the anti-activin-treated mice compared with the IgG-treated controls (Figure 4G, H). Some of the *CXCL10* expression coincided with αSMA-staining suggesting it came from CAFs (Figure 4G), although analysis of the scRNA-seq data indicated that macrophages and neutrophils were also responsible for the enhanced production of *CXCL10* in the tumours (Figure S4E). Furthermore, co-staining of tumours with CD8 and *CXCL10* revealed that clusters of CD8^+^ T cells were indeed observed around the patches of *CXCL10* staining (Figure S4F).

In summary, treatment of mice with either anti-activin antibody reduces the collagen-rich desmoplasia and promotes infiltration of cytotoxic T cells in an orthotopic breast cancer model.

### Anti-activin antibody treatment results in an increase in tumour size in breast cancer models

Given the switch to a more immune hot phenotype and the reduction in collagen that we observed with the treatment of mice with anti-activin antibodies in the breast cancer model, we expected that we might observe signs of tumour regression. However, in this model, we found that the tumours grew to a slightly larger size when the mice were treated with the anti-activin antibodies compared to the IgG control (Figure 5A; S5A). We observed the same effect when using NSG mice, which lack functional/mature T, B, and NK cells ^58^, suggesting that the adaptive immune system was not responsible (Figure 5B; S5B).

**Figure 5.**
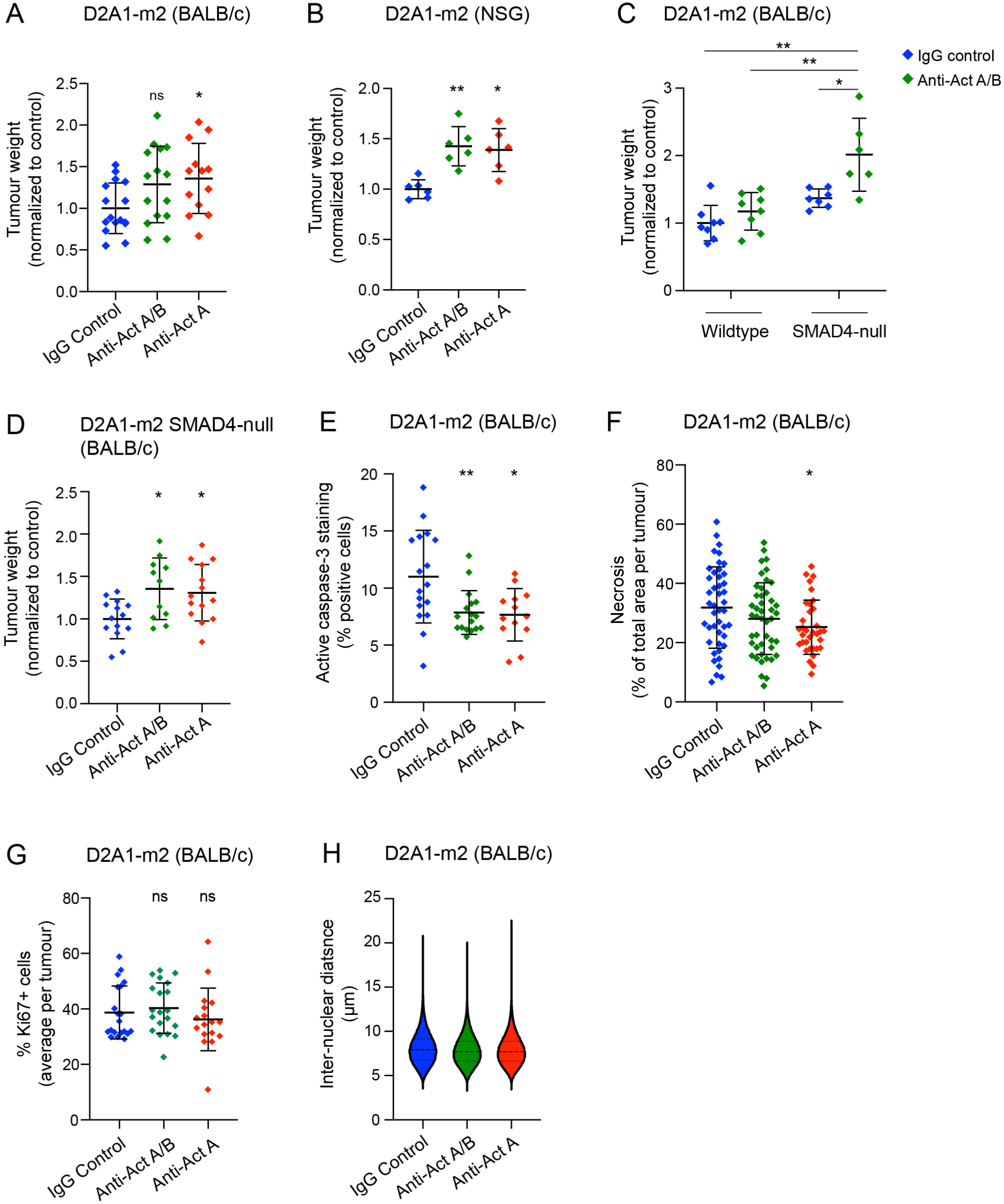
Treatment of mice with the anti-activin antibodies leads to an increase in tumour size as measured by tumour weights. (**A**) Final tumour weights in the D2A1-m2 mouse breast cancer model (D2A1-m2 injected into the mammary fat pad of BALB/c mice). Mice were treated with IgG control, anti-activin A/B or anti-activin A antibodies. Means ± SD are shown. ns, not significant; *, p <0.05. (**B**) As for (A) but NSG mice were used instead of BALB/c. *, p <0.05; **, p <0.01. (**C**) As for (A) but a direct comparison was performed between wildtype D2A1-m2 cells and SMAD4-null D2A1-m2 cells. Mice either treated with IgG control or anti-activin A/B antibodies. *, p <0.05; **, p <0.01. (**D**) As for (A) but SMAD4-null D2A1-m2 cells were used instead of wildtype D2A1-m2 cells. *, p <0.05. (**E**) Quantification of active caspase-3 staining in tumours from BALB/c mice injected with D2A1-m2 cells treated with IgG control (n=17), anti-activin A/B (n=17) or anti-activin A antibodies (n=13). Data are represented as % of total positive cells per tumour. Means ± SD are shown. *, p <0.05; **, p <0.01. (**F**) Quantification of necrotic area in tumours from BALB/c mice injected with D2A1-m2 cells treated with IgG control (n=44), anti-activin A/B (n=45) or anti-activin A antibodies (n=34). Data are represented as % of total necrotic area, per tumour. Means ± SD are shown. *, p <0.05. (**G**) Quantification of Ki67-positive cells in tumours from BALB/c mice injected with D2A1-m2 cells treated with IgG control (n=20), anti-activin A/B (n=20) or anti-activin A antibodies (n=17). Data are represented as % of total positive cells per tumour. Means ± SD are shown. ns, not significant. (**H**) Quantification of the internuclear distances in tumours from BALB/c mice injected with D2A1-m2 cells treated with IgG control, anti-activin A/B or anti-activin A antibodies. Six tumours were analysed per condition. Violin plots are shown with medians and quartiles. In A–D each dot represents a single tumour from an individual mouse. In all cases the statistical significance was calculated relative to the IgG control.

Activin A and B are known to inhibit the growth of tumour cells, acting as tumour suppressors ^20^. We therefore speculated that the increase in tumour size detected when activin activity was neutralized could be the result of inhibiting a direct tumour suppressive function of activin A and B on the cancer cells. To test this, we needed to inhibit the ability of the tumour cells to respond to activin and thus chose to delete SMAD4 which is required for activin signalling, using CRISPR/Cas9 ^6^. However, despite the fact that the SMAD4-null tumour cells do not respond to activin signalling, the anti-activin therapy still enhanced their growth, suggesting in this case its effect may be mediated by targeting the tumour stroma (Figure 5C, D; S5C, D). The increase in tumour size upon systemic inhibition of activin was a widespread phenomenon, as it was also observed in the MMTV-PyMT mouse model of breast cancer ^47^ and in the oral squamous cell carcinoma model, MOC1 ^59^ (Figure S5E–H).

To investigate the underlying mechanism whereby anti-activin therapy caused the small increase in tumour size, we assessed expression of the apoptotic marker, active caspase-3, as well as measuring levels of necrosis, cell proliferation by Ki67 staining, and investigating cell density by measuring internuclear distances (Figure 5E–H). Of these, the biggest effect was a decrease in active caspase-3 staining upon treatment with anti-activin A/B or anti-activin A antibodies, suggesting that inhibiting activin signalling promoted cell survival in these tumours.

### ICIs further increase the ability of anti-activin antibodies to promote cytotoxic T cell infiltration in tumours

The increase in levels of CXCL10 in the TME and increase in CD4^+^ and CD8^+^-positive T cells in tumours from mice treated with the anti-activin therapy prompted us to investigate whether we could achieve an anti-tumour effect by combining the anti-activin antibodies with ICIs, in particular, anti-PD-L1 antibodies ^4,34^. In preclinical models, inhibition of TGF-β, which also promotes infiltration of cytotoxic T cells into the tumours has been shown to synergize with anti-PD-L1 antibody therapy in the EMT6 mouse mammary carcinoma model to induce further T cell infiltration and promote tumour regression ^17^. Similar synergistic effects of anti-TGF-β therapy and ICIs have been reported in other tumour models ^16,60^. This combination therapy has also reduced metastasis in a mouse model of colon cancer ^4,18^.

We noted that the T cells in the tumours from the activin-therapy treated mice showed markers of exhaustion including PD-1 (*Pdcd1*) and LAG3 ^61^, which was most marked in the anti-activin A treated tumours (Figure S6A, B). This might explain why the anti-activin therapy alone was not effective and suggested it might synergize with anti-PD-L1 therapy ^62,63^. Furthermore, slightly elevated levels of *Cd274,* which encodes PD-L1, on macrophages and neutrophils in the TME also pointed to an immune suppressive environment of the tumours treated with anti-activins (Figure S6C). However, we found that the combination of anti-activin therapy and anti-PD-L1 therapy was not sufficient to reverse the increase in tumour size observed with anti-activin therapy alone (Figure 6A) but nevertheless resulted in a marked increase in infiltration of CD4^+^ and CD8^+^ T cells above what anti-activin therapy achieved alone (Figure 6B, C). This is consistent with the idea that interaction between PD-1^+^/CD8^+^ T cells and PD-L1^+^ cDCs in tumour draining lymph nodes, can result in increased proliferation of T cells that enter the circulation and migrate to the tumour ^63^.

**Figure 6.**
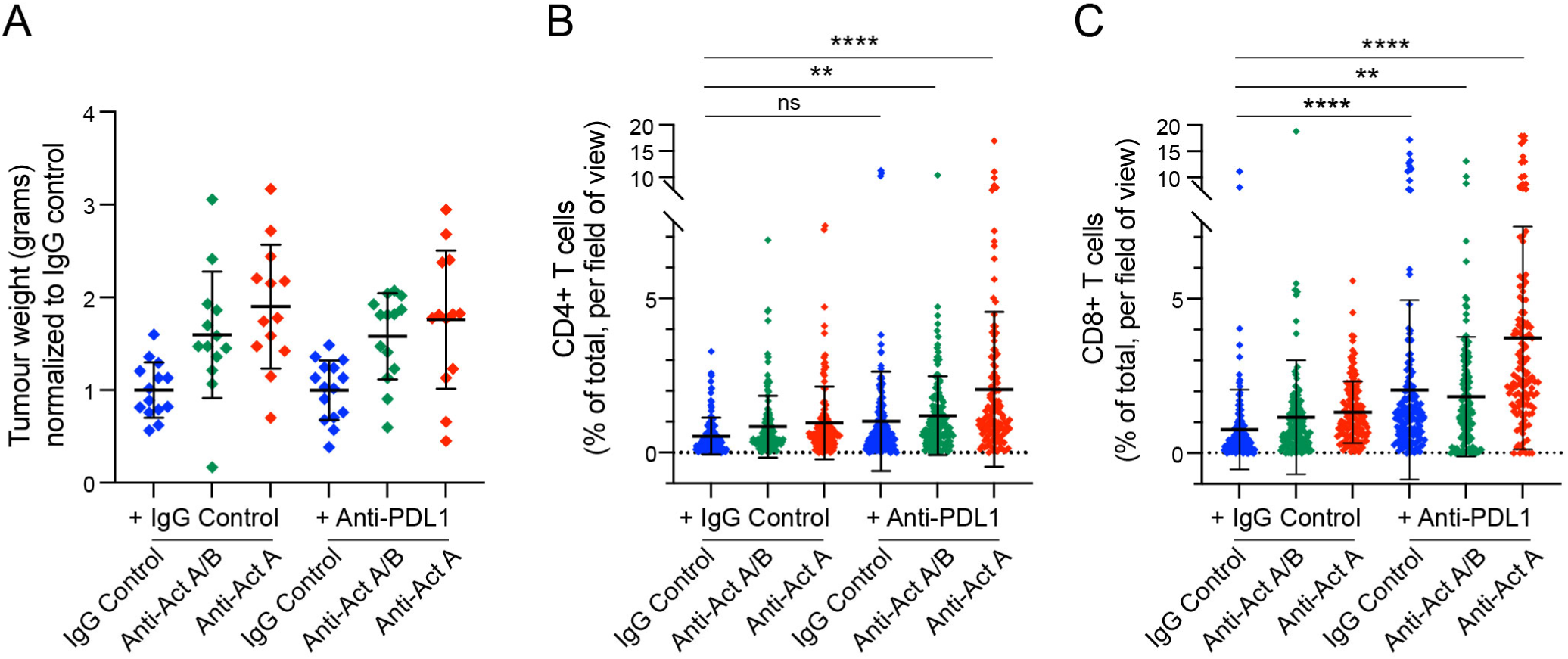
Combining the anti-activin antibodies with immune checkpoint inhibitors or chemotherapy promotes infiltration of CD4^+^- and CD8^+^-positive T cells. (**A**) Final tumour weights in the D2A1-m2 mouse breast cancer model (D2A1-m2 injected into the mammary fat pad of BALB/c mice). Mice were treated with IgG control, anti-activin A/B or anti-activin A antibodies together with a control antibody or with an anti PD-L1 antibody. Tumour weights were normalized to the IgG control group to allow comparison across multiple experiments. There was no statistical significance between the IgG control samples and any of the other samples. Each dot represents a single tumour from an individual mouse. (**B, C**) Quantifications of immune cell infiltration in the mouse breast cancer model (D2A1-m2) from the experiment shown in (A). CD4^+^ (B) and CD8^+^ (C) were analysed in all the tumours. The data are represented as % of total positive cells, per field of view. 20 fields of view were analysed per tumour. ns, not significant; **, p <0.01; ****, p <0.0001.

### Anti-activin treatment in mice with SMAD4-null PDAC tumours promotes survival, but not in tumours containing wildtype SMAD4

Having demonstrated some beneficial effects of anti-activin treatment in the orthotopic breast cancer model, but with no sign of tumour regression, we moved to genetically modified mouse models of PDAC. These mouse tumours also exhibit high levels of *Inhba* and *Inhbb* expression (Figure S1D) and in humans, elevated expression of *INHBA* correlates with poor prognosis (Figure 1A). Importantly, human PDAC is the cancer that most commonly exhibits mutation/deletion of SMAD4 or mutation/deletion of the activin type I and type II receptors ^25–27,64^. This suggests clonal selection ^65,66^ as the tumour evolves for tumour cells that can rapidly grow in an environment rich in activin and/or TGF-β ligands that require SMAD4 for signalling. Moreover, patients with SMAD4 mutations/deletions have a worse prognosis than those with intact SMAD4, suggesting that these tumours are more aggressive ^67^ and confirming the tumour suppressive effect of TGF-β/activins, likely mediated via the tumour cells themselves.

For a wildtype SMAD4 model, we used the genetically modified KPC mouse model (LSL-*Kras*^G12D/+^; *Trp53*^flox/+^; Pdx1-Cre) ^45^. Anti-activin antibodies or IgG control antibodies were injected into the mice as soon as they had developed palpable tumours. These tumours grew very slowly over a period of 6–8 months. We found no effect on mouse survival for the anti-activin A/B antibody compared with IgG control but discovered that the anti-activin A antibody significantly decreased survival (Figure 7A). This is consistent with activin A having a dominant tumour suppressive effect in this model. Interestingly, as in the breast models, we noted an increase in CD4^+^ and CD8^+^ T cells into the tumours after treatment with both anti-activin antibodies, suggesting a possible additional tumour promoting effect of activin mediated via the stroma (Figure S7A, B).

**Figure 7.**
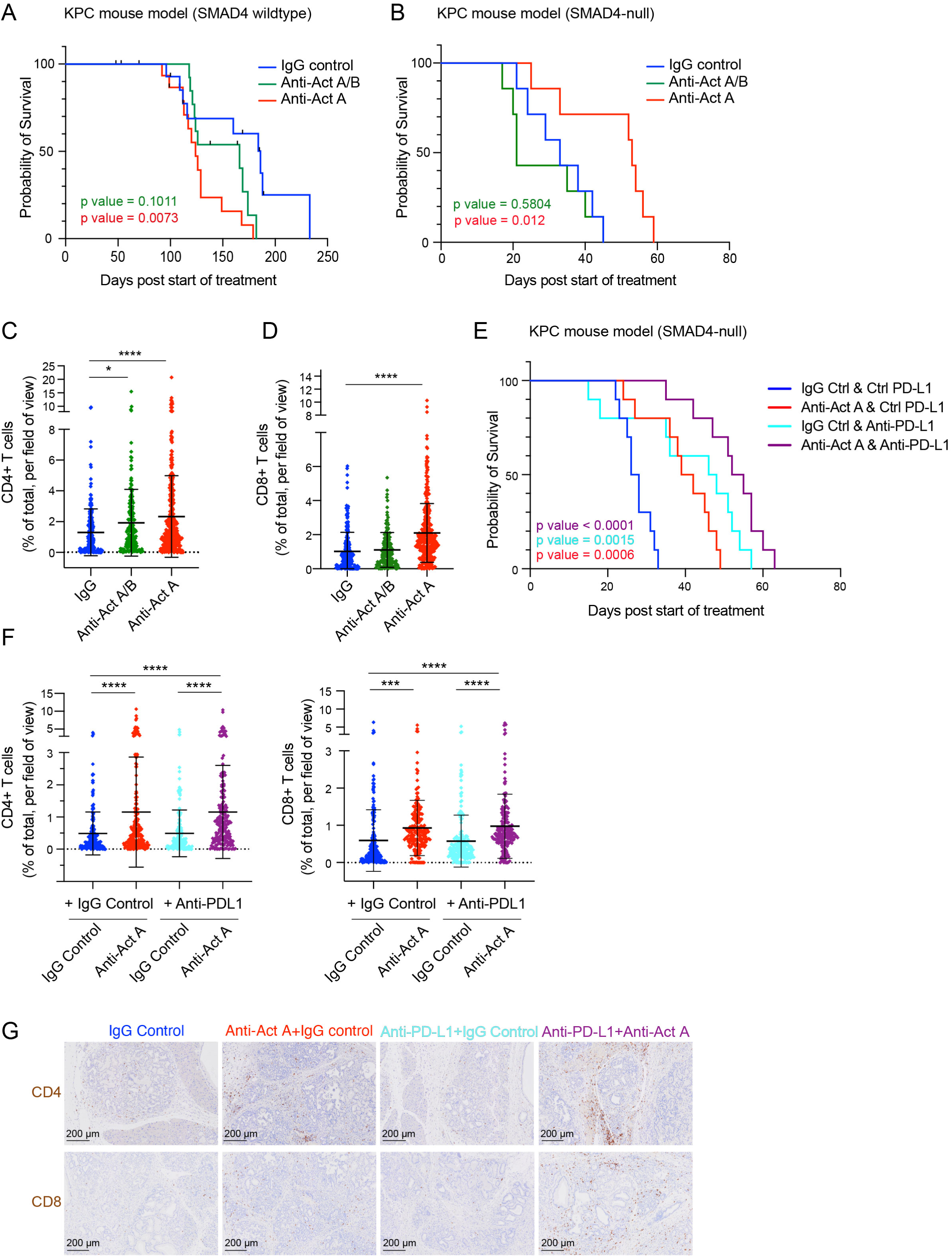
The effects of anti-activin treatment in mouse models of PDAC depends on the status of SMAD4. (**A**) Kaplan Meier survival plots demonstrating overall survival in the wildtype SMAD4 PDAC mouse model (KPC), where mice were treated with IgG control, anti-activin A/B or anti-activin A antibodies. p values are indicated for the differences in the anti-activin A/B (green) or anti-activin A (red) treatments relative to the IgG control. (**B**) Kaplan Meier survival plots for SMAD4-null KPC PDAC mouse model where mice were treated with IgG control antibody, anti-Activin A/B antibody or anti-Activin A antibody. P values are indicated for the differences between the anti-activin A/B (green) or anti-activin A (red) treatments relative to the IgG control. (**C, D**) Quantifications of immune cell infiltration in the tumours in the SMAD4-null KPC mice treated with IgG control, anti-activin A/B or anti-activin A antibodies. CD4^+^ (C) and CD8^+^ (D) cells were detected by immunostaining and quantified in 8 tumours per condition as percentage of cells of interest per field of view. The whole tumour was quantified. 40 fields of view were analysed per tumour. *, p <0.05; **, p <0.01; ***, p <0.001; ****, p<0.0001. (**E**) Kaplan Meier survival plots demonstrating overall survival in the SMAD4-null PDAC mouse model, where mice were treated with IgG control, anti-activin A antibodies with or without anti-PD-L1. p values are indicated for the differences in the anti-activin A (red) or anti-PD-L1 (turquoise) or anti-activin A and anti-PD-L1 treatments (purple) relative to the IgG control. (**F**) Quantifications of immune cell infiltration (CD4^+^ T cells are shown on the left and CD8^+^ T cells are shown on the right) in pancreatic tumours from the SMAD4-null KPC mice treated with IgG control, anti-activin A antibody, anti-PD-L1 antibody or the combination of anti-activin A and anti-PD-L1 antibodies. 20 fields of view are shown for 5 tumours for each condition. Means and SDs are shown. ***, p <0.001; ****, p <0.0001. (**G**) Representative images showing CD4^+^ (upper row) and CD8^+^ cell infiltration in tumours from SMAD4-null KPC mice treated with IgG control, anti-activin A antibody, anti-PD-L1 antibody or the combination of anti-activin A and anti-PD-L1 antibodies as indicated. Tumours were stained with H&E and for the CD4 or CD8 (brown) as indicated. Scale bar, 200 µm.

To determine whether we could uncouple activin’s tumour suppressive and tumour promoting effects, we generated a SMAD4-null PDAC model (*Smad4*^flox/flox^; LSL-*Kras*^G12D/+^; LSL-*Trp53*^R172H/+^; *Ptf1a/p48*-Cre). In this model, the tumour cells cannot respond to activin A/B, owing the lack of SMAD4, but all the cells of the stroma that contain activin receptors will be competent to respond to these ligands, as their SMAD4 is intact. Opposite of what we observed when the tumour cells contained wildtype SMAD4, in SMAD4-null model we observed a substantial increase in survival for mice treated with the anti-activin A antibody compared to those treated with the lower affinity anti-activin A/B antibody, or the IgG control (Figure 7B). This demonstrated that by making the tumour cells null for SMAD4, we had indeed succeeded in removing the tumour suppressive effects of activin. As with the other models, we observed an increase in CD4^+^ and CD8^+^ T cells in the anti-activin treated tumours, which was more profound for those treated with anti-activin A compared to anti-activin A/B (Figure 7C, D). Furthermore, we observed a more robust depletion of pSMAD2 levels in tumours from mice treated with the anti-activin A antibody compared with the anti-activin A/B antibody, consistent with the in vivo effects of these antibodies on mouse survival (Figure S7D, E).

Finally, given the observation that treatment with the anti-activin A antibody of the SMAD4-null PDAC mice promoted survival, and our previous finding in the breast cancer model that additional treatment with anti-PD-L1 antibody further increased cytotoxic T cell infiltration into the breast tumours, we asked whether we could further promote mouse survival by a combined treatment of anti-activin A therapy with an ICI. Indeed, we could demonstrate an additional survival benefit of the combined treatment, compared with either treatment alone (Figure 7E). We saw an increased infiltration of cytotoxic T cells in the presence of both therapies compared with IgG treatment or anti-PD-L1 treatment alone, although no obvious significant increase over that observed with the anti-activin A alone (Figure 7F, G).

## Discussion

### Activin has both tumour suppressive and tumour promoting activities in breast cancer and PDAC

Here we have demonstrated a functional role for activin signalling in mouse models of breast cancer and PDAC and see evidence for both tumour promoting and tumour suppressive functions of activin. In support of a tumour promoting role, we find in both models that inhibiting activin A and/or activin A and B results in increased infiltration of CD4^+^ and CD8^+^ T cells into the core of the tumour, converting the tumours from an immune excluded phenotype to a more immune infiltrated phenotype. In the breast cancer model, we have correlated this with an upregulation of the chemokine CXCL10 expression following activin inhibition. This has also recently been reported in mouse models of melanoma ^34^. CXCL10 is well known for being responsible for attracting CD8^+^ T cells in different cancer contexts ^56,68–70^, and our scRNA-seq and fluorescent in situ data suggest that the source of the CXCL10 is predominantly macrophages and CAFs, where its expression must normally be repressed by activin. Infiltration of T cells into the tumours may also be a result of a breakdown of the stromal fortress around the tumour, as inhibiting activin signalling results in a loss of collagen deposition in the breast cancer model, which would make the tumour more porous.

Consistent with an additional tumour suppressive role for activin, we have shown that in breast tumours containing wildtype SMAD4, neutralization of activin results in a small increase in tumour size. This appears to result from a decrease in apoptosis of tumour cells, as opposed to an increase in proliferation. We hypothesized that this could be due to a direct effect of activin signalling in the tumour cells, mediated via the canonical activin pathway via SMAD4 ^6^. However, deletion of SMAD4 in the D2A1-m2 breast cancer cells did not abolish the small increase in tumour size that we detected upon inhibition of activin signalling suggesting that it was not mediated by a direct effect on the tumour cells themselves. Instead, the results suggest that inhibiting activin signalling promotes expression of a factor in a stromal cell type that may promote survival of the tumour cells.

In contrast to breast cancer, where loss of SMAD4 is generally not detected in the human disease ^71^, our results in the PDAC model indicate that activin has a demonstrable tumour suppressive effect on the tumour cells themselves, consistent with the frequent deletion of SMAD4, or acquisition of loss-of-function mutations in SMAD4 seen in human PDAC ^8^. Strikingly, in this model, we were able to disentangle the tumour suppressive and tumour promoting effects of activin. We found that inhibition of activin A signalling in the mouse KPC model where the pancreatic epithelial cells contain wildtype SMAD4 resulted in poorer survival, whereas in the SMAD4-null KPC model where SMAD4 was deleted in the pancreatic epithelial cells, inhibition of activin A signalling resulted in an increase in mouse survival. This suggests that activin expressed in the TME normally constrains tumour growth, and loss of SMAD4 releases these constraints, allowing activin to act as a tumour promoter via its activity on stromal cells. We hypothesize that one of the main tumour-promoting functions of activin in this context is immune suppression, excluding cytotoxic T cells from the tumour. Consistent with this, in the SMAD4-null PDAC model we saw an additive effect of anti-PD-L1 therapy and anti-activin therapy, as measured by mouse survival.

### Immune checkpoint inhibitors in PDAC and breast cancer

Given activin’s well-described immune suppressive effects, similar to those described for TGF-β ^3,24,72^, and our observation that neutralizing activin activity promoted infiltration of cytotoxic T cells into the tumour core, we predicted that anti-activin therapy would potentially synergize with anti-PD-L1 therapy, as has been observed for TGF-β ^17,18^. In the breast cancer model, we observed a further increase of CD4^+^ and CD8^+^ T cells into the tumour, but no obvious tumour regression. There could be many reasons for this. In these experiments, the anti-activin therapy increases the size of the tumours, meaning that the therapeutic window is short. Thus, the effects of the anti-PD-L1 therapy may not have been realised before the mice needed to be culled due to tumour burden. Another possibility is that the numbers of T cells accumulating may be insufficient for effective killing of tumour cells. Alternatively, the tumours may not have a sufficiently high mutation burden to produce recognized neoantigens ^73^, or the infiltrated T cells may have compromised cytotoxic activity ^62,63^. Furthermore, the presence of other immune suppressive cells in the tumours like tumour-associated macrophages or T_reg_ cells or myeloid-derived suppressor cells can also confer resistance to ICIs ^3,73,74^.

In contrast, in the SMAD4-null PDAC model, we did observe an improved response to the combined anti-activin A and anti-PD-L1 therapy over anti-activin treatment alone. Further work will now be required to understand the mechanism whereby anti-activin therapy has a survival benefit in this model, and how this is augmented by ICIs.

### Potential of anti-activin therapy for cancer

Here we have compared the behaviour of two anti-activin antibodies in our mouse models – a dual specificity anti-activin A/B antibody that blocks the type II receptor binding site and a higher affinity anti-activin A antibody that likely blocks the type I receptor binding site, both in the same IgG1 format. Because of the difference in their affinities for activin A, we have been unable to rigorously define the relative roles of the two activin isoforms, which we find to be expressed in different cell types in the tumours (both human and mouse). We have observed that the anti-activin A antibody is more effective in vivo both for inhibiting activin signalling via pSMAD2, and also in promoting phenotypic effects. This is likely due to its much higher affinity for activin A than our dual specificity anti-activin A/B antibody. In future work it will be important to investigate whether there is any added advantage of targeting activin B. From our RNAScope analyses, activin B appears more broadly expressed in breast and PDAC tumours than activin A, but at lower levels, as evidenced by our scRNA-seq analyses. Activin B also exhibits a distinct receptor repertoire, being able to signal through ACVR1C as well as ACVR1B, whilst activin A is specific for ACVR1B, which might influence their functions in the tumour ^75,76^.

In terms of the potential of anti-activin therapy for solid cancers, our results indicate that patients with PDAC tumours that have lost SMAD4 or acquired loss-of-function SMAD4 mutations would be the most likely to benefit. Moreover, these patients have lower survival than those whose tumours contain wildtype SMAD4 and also have tumours more likely to metastasize ^67,77^. In our mouse models, it was evident that by deleting SMAD4 specifically in the tumour cells we could specifically remove the tumour suppressive effects of activin, whilst maintaining the tumour promoting effects. These tumour promoting functions of activin are evidently mediated via the SMAD4-containing stromal cells, and we can inhibit these functions by neutralizing the activin ligand in the TME.

Anti-activin therapy is expected to be well tolerated in humans and mice. We have found no evidence of side effects in the mice, in agreement with previous reports using an activin A neutralizing antibody in mouse models of fibrodysplasia ossificans progressiva (FOP) ^78^ and use of the anti-activin A antibody garetosmab in humans in non-cancer settings revealed it to be well tolerated with an acceptable safety profile ^38^. Furthermore, other reagents that inhibit activin signalling like galunisertib or vactosertib, that inhibit the kinase activity of the activin and TGF-β type I receptors, are also well tolerated ^3,79,80^.

### Combination therapies for PDAC

As well as combining anti-activin therapy with ICIs in SMAD4-null PDAC, it is important to consider the additional role of the TGF-β ligands. We have shown a reduction of pSMAD2 in both the breast cancer model and PDAC models upon anti-activin therapy, but there are still substantial levels of pSMAD2 in the tumours, suggesting either that the anti-activin antibodies are not potent enough to neutralize all the activin, or that other ligands that activate SMAD2-dependent signalling are also contributing. The obvious candidates would be the TGF-β ligands themselves and in the future, we will investigate the additional benefit of combining anti-activin treatment with the 1D11 antibody that neutralizes all three TGF-β ligands ^81^. Anti-TGF-β therapy has the potential to increase further the response of CD8^+^ T cells to CXCL10, as it has been shown in a colorectal cancer model that the receptor for CXCL10, CXCR3 which is present on CD8^+^ T cells, is negatively regulated by TGF-β. TGFBR1/ALK5-deficient CD8^+^ T cells exhibited increased CXCR3 expression and enhanced migration towards sources of CXCL10 ^68^.

Other combination therapies that could be effective would be anti-activin therapy combined with a chemotherapy such as gemcitabine ^42^ that might increase the access of cytotoxic T cells to the core of the PDAC tumours. Another possibility would be combining anti-activin and anti-TGF-β therapy with radiation therapy. Radiation is known to induce a rapid release of bioactive TGF-β1 from latent complexes and to increase TGF-β1 expression ^82^, as well as increasing activin secretion ^83^. Neutralizing TGF-β and activin has been suggested to reduce regulatory T cell-mediated immunosuppression driven by radiation therapy ^84^, as well as reducing fibrosis and extracellular matrix production ^85^.

## STAR✭METHODS

## RESOURCE AVAILABILITY

### Lead Contact

Further information and requests for resources and reagents should be directed to and will be fulfilled by the lead contact, Caroline Hill.

### Materials Availability

All unique/stable reagents generated in this study are available from the lead contact with a completed materials transfer agreement.

### Data and Code Availability

This paper does not report new datasets or any original code. Any additional information required to reanalyse the data reported in this paper is available from the lead contact upon request. The Cryo-EM map for the activin A–anti-activin A/B Fab complex has been deposited in the Electron Microscopy Data Bank (EMDB), accession number EMD-53342.

## EXPERIMENTAL MODEL AND STUDY PARTICIPANT DETAILS

### Mice

All animal experiments were performed under the Animals (Scientific Procedures) Act 1986 in accordance with UK Home Office license (Project Licenses PP6038402 and PP5869518) and the EU Directive 2010. The Home Office licenses underwent full ethical review and approval by the Francis Crick Institute’s Animal Ethics Committee. The mice were housed in constant temperature, humidity, and pathogen-free controlled environment (25 °C ± 2 °C, 50–60%) cages with a standard 12 h light/ 12 h dark cycle and were allowed access to food and water *ad libitum*. The KPC mouse model was derived by in-crossing the following previously described lines: *Kras*^LSL-G12D/+^ ^86^, Trp53^tm1Brn/+^ (also known as *Trp53^loxP/+^* where loxP sites flank exons 2–10) ^87^ and *Tg(Pdx1-cre)6Tuv* (also referred to as *Pdx1*-Cre) ^88^ as described ^45^. The SMAD4-null PDAC mouse model resulted from in-crossing the following lines: *Kras*^LSL-G12D/+^ (as above), *Trp53*^LSL-R172H/+^ ^89^; *Smad4*^tm1.1Rob/tm1.1Rob^ (also referred to as *Smad4^flox/flox^)* where the first coding exon of *Smad4* is flanked by loxP sites ^90^ and *Ptf1a^tm1.1(cre)Cvw^*(also referred to as *P48*-Cre) ^91^. All strains were genotyped by Transnetyx. NSG (NOD.Cg-*Prkdc*^scid^ *Il2rg*^tm1Wjl^/SzJ) mice were as described ^58^. Wildtype strains used were: BALB/cJ, FvB/NJ and C57BL/6J and were purchased from the Jackson Laboratory.

### Tissue culture cell lines

HaCaT cells were obtained from the Francis Crick Institute Cell Services from stocks deposited in the ICRF Cell Bank by Lional Crawford in 1994. D2A1-m2 cells were provided by Clare Isacke (Institute of Cancer Research, UK) ^46^ and SMAD4 was knocked out in this line using CRISPR/Cas9. The HEK293T CAGA_12_-Luciferase/Renilla cell line was as described ^92^. The CAF1 cell line and normal fibroblasts (NFs) were obtained from Erik Sahai ^48^, as were the MOC1 cell line ^93^. Activin A and B were knocked out in the CAF1 line using CRISPR/Cas9. The MMTV-PyMT primary tumour cells were obtained from transgenic FVB/N mice expressing the Polyoma Middle T-antigen (PyMT) oncogene under the Mouse Mammary Tumor Virus promoter (MMTV-PyMT) ^47^ as previously described ^94^. FreeStyle™ 293-F cells were obtained from the Francis Crick Institute Cell Services.

### Human samples

Anonymised formalin-Fixed Paraffin-Embedded sections of human invasive breast tumours were obtained the Kings Health Partners Cancer Biobank with full ethical approval (REC reference 12/EE/0493 and HTA licence number 12121). Anonymised formalin-Fixed Paraffin-Embedded sections of human pancreatic tumours with matched serum samples were obtained from the Wales Cancer Bank with full ethical approval (Wales REC 3 reference 16/WA/0256 and HTA licence number 12107). Serum from age and gender matched healthy volunteers was obtained from via the Crick healthy volunteers blood donation programme with full consent.

## METHOD DETAILS

### Antibody generation

#### Bead-based selection for Activin scFv binders

The anti-activin A/B antibody was generated by screening phage display libraries. Mature human activin A and *Xenopus laevis* activin B (96.5% identical to human) were expressed in E. coli, refolded from solubilized inclusion bodies and purified to homogeneity as previously described ^95^. Labelling with biotin was performed using EZ-Link™ NHS-PEG4-Biotin (Thermo Scientific, cat# A39259) in a similar way to that described for fluorescently labelled activin A ^95^. To identify activin A/B-specific phage, scFvs were selected by performing three rounds of panning on biotinylated mature activin A and B using phage display libraries ^96,97^ composed of three sub-libraries ^98–100^. Briefly, for each round of selection, phage and Dynabeads™ M-280 Streptavidin (Invitrogen, 11206D) were blocked using 3% milk (Marvel)/PBS. Blocked phage library and beads were co-incubated in the absence of antigen to deplete non-specific phage binders. Subsequently, biotinylated antigens were added, bound phage-antigen complexes captured, and beads washed 5 times with 0.1% Tween-20 (Sigma-Aldrich)/PBS (PBST). Phage were eluted in 10 µg/ml trypsin (BDH) at 37°C. A 5-ml prep of TG1 *E. coli* was grown to mid-log phase and then infected with eluted phage-scFv and plated onto 2TY + ampicillin + glucose (2TYAG) bioassay plates overnight. Colonies were collected into 2TYA media and cultured to mid-log phase before infection with M13 trypsin-cleavable helper phage and overnight selection in 2TY + ampicillin + kanamycin (2TYAK). Phage were recovered by centrifugation at 13,000 rpm for 10 min. The resulting supernatant containing the rescued output was used for the next round of panning. For the first round of panning, 200 nM of biotinylated activin was used, 50 nM was used for the second round and 10 nM was used for the final round.

#### Phage ELISA

After the third round of panning, individual bacterial colonies were picked and cultured in 2TYA overnight and then used to seed fresh cultures which were infected with M13 helper phage once they reached mid-log phase. Cells were pelleted and grown overnight in 2TYAK media, before phage were recovered by centrifugation at 1820 g for 30 mins. Meanwhile, MaxiSorp™ ELISA plates were streptavidin coated overnight and then prepared with 50 µl biotinylated activin A or activin B according to manufacturer’s instructions. ELISA plates and the phage prep were blocked in 3% milk/PBS, before phage were transferred into ELISA plates and incubated at room temperature for 1 h, followed by washing three times with PBST and three times with PBS. Anti-M13 HRP-conjugated antibody (Sino Biological 11973-MM05T-H), 1:5000 diluted, was added to each well and plates incubated for 1 h at room temperature before washing three times with PBST and three times with PBS. Bound antibody was detected using 50 µl TMB substrate (Merck CL07) for 2–5 mins and then 50 µl 0.5 M sulphuric acid was added to stop the reaction. Optical density was measured at 450 nm using an EnVision plate reader (PerkinElmer).

#### Peri-preps and titration ELISAs

Binders from the phage ELISA were further validated by producing bacterial peri-preps. 5-ml bacterial cultures in 2TYAG were established using reserved phage-infected bacterial preps and incubated at 37°C overnight, with 500 µl of this culture then being used to inoculate a further 50-ml culture of 2TYAG. This culture was grown until OD600 reached 0.6-0.8 and then expression of scFvs induced by addition of 1 mM IPTG (Sigma, I6758), with cultures grown overnight at 20°C. The following day, cultures were harvested by centrifugation at high speed for 15 mins, pellets resuspended in 300 µl ice cold TES and 700 µl 10 mM MgSO_4_ added. Preps were centrifuged at high speed for 30 min, and the resulting supernatant collected. These preps were incubated on ELISA plates as for the phage ELISA, with or without multi-point titration. Binders with high affinity and evidence of linear titration were progressed for Sanger sequencing of scFv regions. Primers complementary to the phagemid vector and annealing either side of the scFvs were used to amplify each clone, which was then submitted to an external Sanger sequencing provider (Azenta). The resulting data were then analysed using an internal AstraZeneca proprietary sequence analysis software to identify unique clones.

#### Reformatting and expression

High affinity binders were initially reformatted into pVITRO1-dV-IgG1/lambda as described ^101^. The antibodies were expressed in FreeStyle™ 293-F cells and purified on a Hi-Trap Protein A Column (Pierce), followed by gel filtration using a HILOAD 16/60 SUPERDEX 200 PG (Cytiva) column. These preparations were used for the data shown in Figure 3A–C. For subsequent experiments antibodies were reformatted into pEU1.2 (for the VHs) and pEU4.4 (for the VLams) AZ proprietary vectors and expressed and purified in-house at AstraZeneca as previously described ^102^. The anti-activin A-specific antibody was produced the same way (pEU1.2 for the VH and pEU3.4 for the Vkappa AZ proprietary vectors) using publicly available sequences (https://gsrs.ncats.nih.gov/ginas/app//substances/KR9ZSKO5QE). The NIP228 negative control antibody was provided by AstraZeneca ^103^. Anti-PD-L1 antibodies and relevant negative controls were provided by AstraZeneca ^104^.

### Biolayer interferometry and surface plasmon resonance (SPR)

Biolayer interferometry was carried out using an Octet RED96 instrument (ForteBio) with Octet Kinetic buffer (1X PBS with 0.1% w/v BSA and 0.02% v/v Tween-20). Initial screening was performed by immobilizing purified HEK293T-expressed antibodies at a concentration of 5 µg/ml to anti-human IgG Fc tips (Sartorius). The antibody-bound tips were then probed in activin A or activin B containing solutions at concentrations ranging from 1 to 50 nM. The sensors were regenerated by repeated washing in 10 mM glycine pH 1.7. For verification of the binding to type II receptor binding site, the activin A solution was spiked with the extracellular domain (ECD) of the type II receptor, ACVR2B, produced in *E. coli* by secretion to periplasmic space as fusion protein with Thioredoxin reductase as described previously for ACVR1L and BMPR2 ^105^. Competition was also performed with follistatin-288, purified as previously described ^50^. To test for competition between the anti-activin A/B and anti-activin A antibodies, biotinylated activin A was loaded onto streptavidin sensor tips at 1 µg/ ml. The anti-activin A/B and anti-activin A fragment antibody binding regions (Fabs) were probed consecutively without washing at concentrations of 50 nM. The anti-activin A/B antibody was immobilised on Octet Biosensor anti-human Fc capture (AHC) sensors at a concentration of 5 µg/ml in Octet Kinetic buffer. Dissociation constants were derived from kinetic binding data using Octet BLI system software package using a partial dissociate model.

SPR binding studies were conducted using the BIAcore T200 system (Cytiva) and analyzed with Biacore T200 Analysis Software v3.0. Activin A ^106^, TGF-β3 ^106^ and BMP2 (Peprotech) were immobilized on a CM4 chip (Cytiva) through EDC-NHS activation, followed by injection of 0.5 µM ligand in sodium acetate buffer (pH 4.5) until a maximum increase of 400 response units (RU) was achieved. The chip’s functionality was validated by running a series of dilutions of ligand-specific binders: ACVR2B-ECD ^106^, TGFBR2-ECD ^107^, and Noggin (Peprotech), respectively. All antibody binding experiments were performed in a buffer consisting of 10 mM CHES, 150 mM NaCl, 3 mM EDTA, and 0.01% P20 surfactant at pH 8.0. The surface was regenerated between each injection with a 10-second injection of 0.2 M guanidine hydrochloride at pH 2.0. Experimental sensorgrams were obtained using double referencing with a control surface and four blank buffer injections. Data analysis was performed by fitting the results to a 1:1 kinetic model using the SPR Evaluation Software (Cytiva).

### Expression and purification of the anti-activin A/B Fab complex

Anti-activin A/B heavy and light chains were cloned into Fab constructs in *E. coli* expression vectors pOP3BT (Addgene #112603) with an N-terminal His tag and GB1 fusion and pBAT4 (Addgene #112580), respectively. Anti-activin A/B heavy and light chains and activin A were then expressed insolubly in BL21 (DE3) *E. coli* for 3 h at 37 °C. After solubilization and refolding of activin A for over 7 days as described ^95^, solubilized anti-activin A/B light and heavy chain inclusion bodies were added to the activin A refolding solution for an additional 7 days to allow for the full complex to form. The activin A–anti-activin A/B Fab complex was initially purified by affinity chromatography with a HisTrap Excel column (Cytiva) followed by size-exclusion chromatography with a Superdex 200 Increase 10/300 GL column (Cytiva) in 10 mM HEPES, 200 mM NaCl, 0.5 mM EDTA, pH = 7.5 to obtain a homogeneous sample.

### Cryo-EM sample preparation and data collection

The purified activin A–anti-activin A/B Fab complex was prepared for cryo-EM studies at a concentration of 0.2 mg ml^-1^. Holey copper grids (Quantifoil) with 300 mesh and R1.2/1.3 hole spacing were glow discharged for 60 s at 25 mA and 0.38 mBar. 3 µl of sample was applied, and the grid was blotted for 3 s at 4°C and 95% relative humidity then plunge frozen with liquid ethane by Vitrobot (ThermoFisher). Grids were clipped and screened with a Titan Krios transmission electron microscope at the University of Cambridge Nanoscience Centre before collection of 16,056 movies with pixel size 0.654 Å.

### Cryo-EM data processing

Data processing was conducted with CryoSPARC V4. Movies were motion and CTF corrected with the CryoSPARC patch implementations. Poor quality micrographs were removed with 13,954 movies remaining. 1,000 micrographs were used to generate an initial structure to create 2D templates. Particles were extracted using these templates with a box size of 700 pixels Fourier cropped to 240 pixels from all micrographs, resulting in 1,108,374 particles. Multiple rounds of 2D classification and *ab initio* model generation were performed to determine the morphology of the complex and generate three classes for a heterogeneous refinement (2 “true” classes and 1 “junk” class) of elliptical blobs of the predicted size of the activin A–anti-activin A/B Fab complex. Particles from the “true” class, representing ∼72% of extracted elliptical blobs with noise subtracted (329,495 particles) were used for a final global non-uniform refinement to generate a C1 map with a global masked resolution of 5.40 Å (Figure S8). Resolution of the full complex was significantly limited by the flexibility of activin A.

Models for anti-activin A/B and anti-activin A antibodies with activin A and/or B were generated using the AlphaFold3 server (https://alphafoldserver.com) using two copies of the mature domain of activins, two copies of antibody light chains and two copies of the Fab part of the antibody heavy chains. The models were fitted to cryo-EM density using the Chimera ‘Fit in Map’ tool.

### Cell culture

HaCaT cells, HEK293T CAGA_12_-Luciferase/Renilla, D2A1-m2, D2A1-m2 SMAD4 null and MOC1 were grown in Dulbecco’s Modified Essential Medium (DMEM) (Gibco) with 10% FBS and 1X Penicillin/Streptomycin (Pen/Strep) solution (ThermoFisher Scientific). NF1 and CAF1 cells and their derivatives were grown in DMEM including 1X Insulin-Transferrin-Selenium (ITS), 10% FBS and 1X Pen/Strep. FreeStyle™ 293-F cells were grown in 293 Freestyle media (ThermoFisher) with 50 μg/ml Hygro Gold (Invivogen). MMTV-PyMT cells were grown in DMEM/F12 1:1 (ThermoFisher Scientific), 1X L-Glutamax (ThermoFisher Scientific), 2% FBS, 1% PenStrep, 10 μg/ml insulin (Sigma) and 20 ng/ml EGF (PeproTech). Activin A and activin B-null CAF1 cells were generated using CRISPR/Cas9 using the guides given in Supplementary Table S1. Activin A&B-null clones were identified by Western blotting and confirmed by sequencing (data not shown). Two clones were picked for subsequent experiments. SMAD4-null D2A1-m2 knockout cells were generated using CRISPR/Cas9 as previously described ^108^. The guides are given in Supplementary Table 1. SMAD4-null clones were identified by Western blotting (data not shown). A pool of four individual clones was used for the in vivo experiments.

To prepare conditioned media from CAF1 and NF1s, cells were plated in 10 cm dishes and incubated for 72 h. Media was then harvested and centrifuged at 3000 g in a Centriprep centrifugal filter unit (Millipore) to achieve a four-fold concentration.

Details of recombinant activin A, TGF-β1, follistatin and blocking TGF-β antibody used in the cell-based assays are given in the Supplementary Table 2.

### Western blotting, luciferase assays and qPCR

Whole-cell extracts were prepared as previously described ^92^. Western blots were carried out using standard methods. The list of the antibodies used is shown in Supplementary Table 2. Luciferase assays and qPCR assays were performed as described ^92^ with qPCR primers listed in Supplementary Table 1.

### Contraction assays

For contraction assays, parental CAF1 cells, activin A&B-null CAF1s and NF1 cells were seeded in collagen suspensions (made on ice) comprising 60,000 cells in 100 µl of type I collagen essentially as previously described ^48^. Where appropriate, gel mixes were supplemented with 10 µM of SB-505124 or 20 ng/ml activin A as indicated. Once seeded, gels were left to set for 45 minutes at 37°C. Media was added to gels with or without 10 µM SB-505124 or 20 ng/ml activin A. Gels were imaged immediately and at every subsequent 24-h time point for 72 h with a Leica MZFLIII light microscope. Gel areas were quantified using FIJI software.

### Mouse experiments

#### Orthotopic models and genetically modified mouse models

For the breast cancer D2A1-m2 model, D2A1-m2 cells were washed in PBS and resuspended in 90% growth factor reduced (GFR) Matrigel (Corning, cat# 356231) in PBS. 25,000 cells in 50 μl of GFR Matrigel were injected unilaterally into the lower mammary fat pad of BALB/cJ mice. Isofluorane was used during the injection to anaesthetise the mice. For the MMTV-PyMT breast cancer model, 2 x 10^5^ primary MMTV-PyMT tumour cells were injected into the mammary fat pad of FvB/NJ mice as described above. For the MOC1 model, 1 x 10^6^ MOC1 cells in 100 µl 30% GFR matrigel in PBS were injected subcutaneously into C57BL/6J mice.

The KPC and the SMAD4 KPC PDAC mouse models were generated by in-crossing genetically modified mice as described above. The target genotype for the KPC mice was Pdx1-Cre; *Trp53^loxP/+^*; *Kras*^LSL-G12D/+^, and the target genotype for the SMAD4 KPC mice was p48-Cre^+/-^; Smad4^flox/flox^; Kras^LSL-G12D/+^; Trp53^LSL-R172H/+^.

#### Therapeutic treatments with antibodies

After confirmation of a palpable tumour (D2A1-m2, MMTV-PyMT breast cancer models or MOC1 oral squamous cell carcinoma model) or identification of a mixed cystic lesion or tumour measuring 0.3 x 0.3 cm on ultrasound imaging (PDAC models), 400 μg of anti-activin A/B, or anti-activin A or IgG control antibodies freshly diluted in PBS (20 mg/kg) were injected intraperitoneally (IP) twice a week. When tumours reached endpoint, mice were culled by cervical dislocation and tumours removed. Where anti-PD-L1 antibodies and relevant controls were used, these were injected at 10 mg/kg. For the first two weeks of treatment a mouse anti-PD-L1 (D265A clone 80) or its matched control ^104^ was used. As an anti-drug response can occur, after 2 weeks of treatment with murine antibodies, mice were swapped onto a rat anti-PD-L1 antibody (clone 10F.9G2) or a rat IgG2b control for 1 week of further treatment, after which they reverted onto the murine anti-PD-L1 antibody or control, in a continuous cycle.

In the orthotopic models, tumour size was used as the primary endpoint. Tumour volumes were measured using callipers for mammary fat pad or the subcutaneous tumours. Mice were culled either when tumours reached 1.8 cm^3^ (corresponding to width x length: ∼1.5 cm x 1.5 cm), or when other humane endpoints were met. All mice in the experiment were culled at the same timepoint. For murine PDAC models, survival was used as the primary endpoint, and mice were culled if they demonstrated various signs of ill health, such as piloerection, significant weight loss, swollen/sunken abdomen or huddled posture. Tumour size was measured during the experiment using ultrasound. Tumour weights were measured at the end of the experiment once tumours had been excised.

#### Blood collection and plasma preparation from mice

Blood was collected from mice either by cardiac puncture at the end of the experiment, followed by culling by cervical dislocation or from the saphenous vein (<50 µl per sample) at intervals during in the experiment (spaced 2 or 3 days apart).

Where the volume was sufficient, the collected blood samples were placed in lithium EDTA tubes. Samples were centrifuged at 1,500 g for 15 min at 4°C. The plasma was collected and stored at -80°C. For very small blood volumes heparin-coated microvettes were used to collect the blood, which was centrifuged at 3000 rpm for 15 min at 4°C, before collecting the plasma and freezing at -80°C.

### Assaying levels of activin A, activin B and IgG in the circulation of mice and humans

Human and mouse activin A and B ELISAs were obtained from Ansh Lab (human Activin B, AL-150, mouse Activin B AL-156) and R&D (human/mouse Activin A, DAC00B), and used according to the manufacturer’s instructions. Levels of injected human antibodies in mouse plasma from FvB mice injected IP with 400 µg of antibody (day 0). Blood samples were taken at day 1, 4 and 7. Plasma antibody levels were assayed by ELISA using a human IgG1 ELISA kit (Thermo Scientific, EHIGG1) according to the manufacturer’s instructions.

### Immunohistochemistry and immunofluorescence of FFPE tumour sections

Tumours were fixed overnight in 10% neutral buffered formalin and subsequently stored in 70% ethanol. Samples were then embedded in paraffin and sectioned. Tissue sections were baked for 1 h at 60°C, deparaffinised and rehydrated using standard methods. Heat-mediated antigen retrieval in citrate buffer pH 6.0 or Tris EDTA buffer pH 9.0 was performed, followed by blocking of endogenous peroxidase in 1.6% H_2_O_2_ in PBS for 10 min. Slides were washed in dH_2_0, and background was blocked with a solution of 4% BSA in PBS. For a list of the primary antibodies used, see Supplementary Table 2. Secondary antibodies were HRP conjugated. Where the signal was weak, biotin-conjugated secondary antibodies were used, diluted in 5% BSA in PBS and incubated for 45 min. All washing was performed for 5 min, twice with PBS, and once in PBS with 0.05% Tween-20. The VECTASTAIN ABC-HRP kit and DAB chromogen (Vector Laboratories) were used according to manufacturer’s instructions, and the slides were subsequently counterstained with haematoxylin and mounted. Slides were imaged at 10X with the Zeiss AxioScan slide scanner with Zen Blue software and quantitated using Qupath.

For immunostaining for pSMAD2, sections were prepared as above but endogenous peroxidase was blocked in 3% H_2_O_2_ in methanol for 30 min. All washing was done three times for 5 min in TBS with 0.05% Tween-20 (TBS-T). The primary antibody was incubated overnight at 4°C, followed by 1 h incubation with biotinylated donkey-anti rabbit (Jackson ImmunoResearch). The TSA biotin system (Akyoa Biosciences, NEL749A001KT) was used to amplify the pSMAD2 signal, according to the manufacturer’s instructions. Briefly, slides were incubated for 30 min with Streptavidin-HRP diluted in 4% BSA in TBS-T, washed, and incubated for 8 min with biotinylated tyramide diluted 1:50 in amplification diluent. After another wash, slides were incubated for 30 min with Streptavidin Alexa Fluor-488 (Molecular Probes) and DAPI in 4% BSA in TBS-T. Slides were then incubated for 15 min in a solution of 0.1% Sudan Black in ethanol, washed 3 times for 10 min in TBS-T and mounted. For αSMA immunofluorescence, sections were treated as above, except that the tyramide amplification step was not required and a donkey anti-mouse Alexa Fluor-594 secondary antibody (Invitrogen) diluted in 4% BSA in TBS-T was used instead. Imaging was performed using the Olympus VS120 Slide Scanner or the Zeiss Axio Scan Z1 with Zen Blue software and quantitated using Fiji or Qupath.

### Multiplex RNA in situ hybridization (RNAscope™)

Tissue sections were baked for 1 h at 60°C, deparaffinised and rehydrated using standard methods. The RNAscope Multiplex Fluorescent Reagent kit v2 (ACD Cat. No. 323100) was used according to the manufacturer’s instructions. Briefly, sections were pretreated with the H_2_O_2_ solution, boiled at 100°C for 15 min in the target retrieval solution, and incubated with the Protease Plus reagent. A combination of C1 and C2 were diluted as per instructions and hybridized for 2 h at 40°C in the HybEZ oven. For a list of probes see Supplementary Table 2. A series of amplification steps was performed, followed by channel specific HRP incubation steps and Opal dyes (Akoya Bioscience) for each probe used. If the RNAScope was multiplexed with immunofluorescence, the immunofluorescence staining was then performed as described above. Slides were mounted and imaging was performed on a Zeiss Invert 880 NLO confocal microscope and ZEN platform, with a Plan-Apochromat 20X/0.8 objective.

### Collagen staining

Tissue sections were deparaffinised and rehydrated using standard methods and stained for 10 min in Weigert’s working haematoxylin solution, a mix of solutions A and B (Sigma-Aldrich, HT107-500ML and HT109-500ML respectively). Sections were then washed in tap water for 5 min, rinsed in dH_2_O and stained in Picrosirius red solution (Abcam, ab150681) for 1 h. After two quick washes in acidified water, slides were dehydrated in absolute ethanol and mounted.

### Preparation of single cells for scRNA-seq

To obtain single cell preparations of D2A1-m2 tumours from mice treated with IgG, anti-activin A/B or anti Activin A, the mice were culled and tissues collected. Tumours were transferred to ice cold PBS, then to a 10-cm cell culture dish and minced using scissors and a disposable scalpel, to form a paste-like consistency. The tissue was placed in a gentleMACS™ C Tube (Miltenyi Biotec, cat# 130-093-237) with 10 ml of digestion buffer (DMEM with 2% FBS and Collagenase D at 0.4 mg/ml, Dispase II at 0.6 mg/ml and DNAase I at 40 µg/ml). The tissue was digested on the gentleMACS™ Dissociator (Miltenyi Biotec) with heating blocks installed. The digestion mix was then passed through a 100 µm strainer and washed through with DMEM/2% FBS to a final volume of 50 ml. This was centrifuged at 1250 rpm for 10 minutes at 4°C and the supernatant discarded. The pellet was resuspended in 1 ml of 1x RBC Lysis Buffer (Biolegend, cat# 420301) for 3 min at room temperature. This was quenched with 50 ml of Hanks Balanced Salt Solution (HBSS) with 10% FBS and centrifuged as previously. The pellet was resuspended in 5 ml of DMEM/2% FBS and passed through a 70 µm strainer. The suspension was stained with DAPI (0.2 mg/ml) 1:500 for 10 min in the dark on ice. The tubes were centrifuged as previously and resuspended in 1 ml DMEM/2% FBS and filtered into soft polyprolene FACS tubes. Cells were sorted using DAPI positivity as an exclusion criterion to obtain viable cells, which were collected into HBSS containing 0.4% BSA.

### Library preparation and sequencing

For the bulk RNA-seq experiment comparing the transcriptomes of growing parental CAF1 cells with two clones of Activin A/B-null CAF1 cells, total RNA was extracted using TRIzol further purified using the purified using the RNeasy kit (QIAGEN) as per the manufacturers’ instructions. RNA quality was assessed using a Bioanalyzer (Agilent, California, USA). Libraries were prepared using polyA KAPA mRNA Hyper Prep kit (KAPA Biosystems). Single-end reads were generated using the Illumina HiSeq 2500 platform.

For the sc-RNA-seq experiment on D2A1-m2 tumours from IgG, anti-Activin A/B or anti-Activin A-treated mice, the concentration and viability of the single cell suspension, generated as described above was measured using acridine orange and propidium iodide in a Luna-FX7 Automatic Cell Counter. Approximately 5000-20,000 cells were loaded on Chromium Chip and partitioned in nanolitre scale droplets using the Chromium Controller and Chromium Next GEM Single Cell Reagents (CG000315 Chromium Single Cell 3’ Reagent Kits User Guide (v3.1 - Dual Index). Within each droplet the cells were lysed, and the RNA was reverse transcribed. The resulting cDNA within a droplet shared the same cell barcode. Illumina compatible libraries were generated from the cDNA using Chromium Next GEM Single Cell library reagents in accordance with the manufacturer’s instructions (10x Genomics, CG000315 Chromium Single Cell 3’ Reagent Kits User Guide (v3.1 - Dual Index)), with 12 cycles of PCR amplification. Final libraries were QC’d using the Agilent TapeStation and sequenced using the Illumina NovaSeq 6000 with sequencing read configuration: 28-10-10-90.

### Bioinformatics

For the bulk RNA-seq experiment on CAFs, raw reads were quality and adapter trimmed using cutadapt-1.9.1 ^109^ prior to alignment. Reads were then aligned and quantified using RSEM-1.3.0/STAR-2.5.2 ^110,111^ against the mouse genome GRCm38 and annotation release 89, both from Ensembl. TPM (Transcripts Per Kilobase Million) values were also generated using RSEM/STAR. Differential gene expression analysis was performed in R-3.6.1 ^112^ using the DESeq2 ^113^ package (version 1.24.0) with two different contrasts: 1) pairwise comparison between each activin A/B null condition to the parental control; 2) comparison between the combined activin A/B null and the parental conditions. Differential genes were selected using a 0.05 false-discovery rate (FDR) threshold. Normalization and regularized-log transformation (rlog) was applied on raw counts before performing principal component analysis and euclidean distance-based clustering.

Gene ontology (GO) enrichment analysis was performed in R3.6.1 using the clusterProfiler ^114^ package (version 3.12.0) with a p- and q-value cutoff of 0.05. Reactome pathway enrichment analysis was performed in R3.6.1 using the ReactomePA ^115^ package (version 1.28.0) with default parameters. Heatmaps were generated in R-3.6.1 using the ComplexHeatmap package (version 2.0.0) ^116^. For highlighting general changes between Activin A&B null and parental CAFs, average regularized-log transformed counts for each Activin A&B null condition and the parental CAFs were calculated. Subsequently the data were scaled by subtracting the average regularized-log transformed counts of the parental condition from all conditions. Only those genes were visualized that are significantly differentially expressed between the combined Activin A/B null and the parental condition.

For the scRNA-seq analysis of the D2A1-m2 tumours from mice treated with anti-activin A/B, anti-activin A or IgG control the 10X FASTQ-files were aligned with the CellRanger toolkit (10X Genomics, version 5.0.0) toolkit to 10X pre-built mouse reference transcriptome (refdata-gex-mm10-2020-A). Individual samples were assessed using the Seurat R-package (version 4.0.5) ^117^. Cells featuring more than 10% mitochondrial transcripts or less than 200 mRNA features were excluded from the analysis. 30 PCA dimensions were used as basis for integration using the Seurat cca integration method. Clusters were assigned using the Seurat FindClusters function with the cluster parameter set to 0.5. Clusters were assigned based on cell type marker genes, taken from a variety of sources within the literature ^43,118–125^ as well as the human protein Atlas https://www.proteinatlas.org/, PanglaoDB https://panglaodb.se/search.html, <u>and CellMarker</u> http://xteam.xbio.top/CellMarker/index.jsp. Percentages have been calculated based on numbers of cells in each cluster.

For the re-analysis and sub-clustering of the human PDAC scRNA-seq dataset ^43^, count matrices for all samples in this dataset were kindly provided by the Wu lab (24 primary PDAC tumour samples and 11 control pancreas samples). Individual samples were assessed using the Seurat R-package (version 4.0.5) ^117^. Cells featuring more than 20-40% mitochondrial transcripts (depending on the sample) or less than 500 mRNA features were excluded from the analysis. 30 PCA dimensions were used as basis for integration using the Seurat rpca integration method. Clusters were assigned using the Seurat FindClusters function with the cluster parameter set to 0.3. Cell types were assigned based on cluster marker genes. In this first analysis clusters containing fibroblasts were selected for sub-setting. With the fibroblast subset, a new single-cell analysis was conducted with the following parameters: filtering on 20-40% mitochondrial transcripts; 500 mRNA features; 30 PCA dimensions; rpca integration method; Seurat cluster parameter: 0.3. Clusters were assigned based on cell type marker genes as above.

For the re-analysis and sub-clustering of a mouse PDAC single-cell RNA-seq dataset, count matrices for the four KPC viable replicates and the four fibroblast replicates ^125^ were downloaded from GEO, accession number: GSE129455. Samples from this dataset were analysed in separate single-cell analyses: Viable dataset, samples: KPC1_Viable, KPC2_Viable, KPC3_Viable, KPC4_Viable; Fibroblast dataset, samples: KPC1_FibroblastEnriched, KPC2_FibroblastEnriched, KPC3_FibroblastEnriched, KPC4_FibroblastEnriched. For each of the two single-cell analyses the respective individual samples were assessed using the Seurat R-package (version 4.0.5) ^117^. Cells featuring more than 20% mitochondrial transcripts or less than 500 mRNA features were excluded from the analysis. 30 PCA dimensions were used as basis for integration using the Seurat cca integration method. Clusters were assigned using the Seurat FindClusters function with the cluster parameter set to 0.3. Clusters were assigned based on cell type marker genes, as above.

Single-cell differential gene expression analyses were conducted with the glmGamPoi R-package, version 1.1.13 ^126^. MA plots were plotted as average intensity versus log fold change. Volcano plots displayed as log fold change versus adjusted p value.

For the TCGA analyses, RSEM normalized gene expression data was downloaded from TCGA (https://gdac.broadinstitute.org/) for PAAD and BRCA cancer datasets. Healthy and replicates tumour samples were removed. Only one tumour sample per patient was taken forward for downstream analysis. Samples were ranked using the expression of *INHBA*. Patients in the top and bottom quartiles were used to draw Kaplan-Meier plots and significance was tested using the log-rank (Mantel-Cox) test. For the comparison of tumour and normal samples, normalized expression for *INHBA* from matching normal and tumour samples from the same patients (112 of each) was extracted from the BRCA dataset. A paired t-test was used to test significance.

For the Cibersort analysis RSEM normalized expression from tumour samples from the PAAD and BRCA dataset, were used as input to Cibersort ^127^, in order to characterize the cell composition of each sample using 22 immune cells signatures, with 100 permutations. A spearman’s statistic was used to estimate a rank-based measure of association between *INHAA/INHBB* normalized expression and Cibersort’s percentage infiltrate. The level of significance is shown using stars (*, p < 0.05; **, p<0.005; ***, p< 0.0005).

## QUANTIFICATION AND STATISTICAL ANALYSIS

### Quantification of images

All images were quantified in a blinded fashion using QuPath. For image analysis of DAB IHC where multiple fields of view (FOV) were taken within a tumour, annotations were placed randomly throughout the tumour to ensure wide tumour coverage, with either 10, 20 or 40 FOV per tumour, depending on the heterogeneity of the signal. The annotations measured 832 µm width and 666 µm height. Where core and boundary were analysed, core FOVs were placed within the centre of the tumour, avoiding necrotic areas, and boundary FOVs were placed at the outer edge of the tumour, with the entire annotation always within 1 mm of the edge of the tumour. A positive cell or nucleus detection was performed to capture positive cellular staining, with the threshold adjusted using relevant positive controls per experiment. Where entire tumour sections were analysed, a thresholder was used to create a pixel classifier to automate identification of tumour sections. For pSMAD2 immunofluorescence quantification, the positive cell detection settings were set to allow for detection of cells that were mildly, moderately or strongly positive. Measurement map ranges within QuPath were analysed to determine the settings for these 3 categories. The Histoscore was calculated within QuPath using the formula: HistoScore = ((1 × % mildly stained cells) + (2 × % moderately stained cells) + (3 × % strongly stained cells)). To quantitate RNAScope staining, a classifier was used to train QuPath to detect positive, negative and ignore areas based upon the positive and negative controls. The ‘ignore’ areas were necrotic tissue. To measure internuclear distances, a workflow was developed in Qupath to measure the distance between a cell nucleus and its closest neighbouring cell within a tumour.

### Statistical analysis

Unless otherwise stated above, statistical analysis was performed in GraphPad Prism 10. At least three independent experiments were performed for statistical analysis. For comparison between more than two groups, ordinary one-way ANOVA with Dunnett’s multiple comparison test was used or with Tukey’s multiple comparison test when data were normally distributed; alternatively, Kruskal-Wallis with Dunn’s multiple comparison test was used. For pair-wise comparison, two-tailed student’s t-test was used. Data are typically presented as mean ± standard deviation (SD) as indicated in the legends. p values are indicated by: * <0.05, ** < 0.01, *** < 0.001, **** < 0.0001. For assessing significance in Kaplan Meier survival plots, the log-rank (Mantel-Cox) test was used.

## Acknowledgements

We thank Axel Behrens, Jessica Nelson, Karen Vousden, Eric Cheung and Dinis Calado for mouse strains. We thank Erik Sahai for the CAFs, normal fibroblasts and MOC1 cells, Clare Isacke for D2A1-m2 cells and Louise Richardson for the MMTV-PyMT tumour cells. We thank Wenming Wu for giving us access to the human sc-RNAseq dataset from PDAC and control pancreases. We are extremely grateful to staff in the Crick Biological Research Facility for all their help with the mouse experiments and to staff in the Experimental Histopathology, Advanced Light Microscopy, Genomics, Structural Biology and Flow Cytometry facilities at the Crick. We are grateful to the Biophysics Facility at the Department of Biochemistry, Cambridge, for access to instrumentation and support. We thank all the members of the Hill lab past and present for useful discussions as this project was developing and to Simon Barry, Gerard Evan, Lutz Jermutus, Ilaria Malanchi, Anjana Ramdas Nair and Scott Wilcockson for very helpful comments on the manuscript. This work was supported by the Francis Crick Institute, which receives its core funding from Cancer Research UK (CC2021), the UK Medical Research Council (CC2021), and the Wellcome Trust (CC2021), by I2I funding from the Francis Crick Institute and by funding from AstraZeneca. GP is supported by a PhD studentship from The Gates Cambridge Trust.

Note that the pancreatic tumour samples and plasma were obtained from the Wales Cancer Bank which is funded by the Welsh Government and Cancer Research Wales. Other investigators may have received specimens from the same subjects.

---

**Figure S1.**
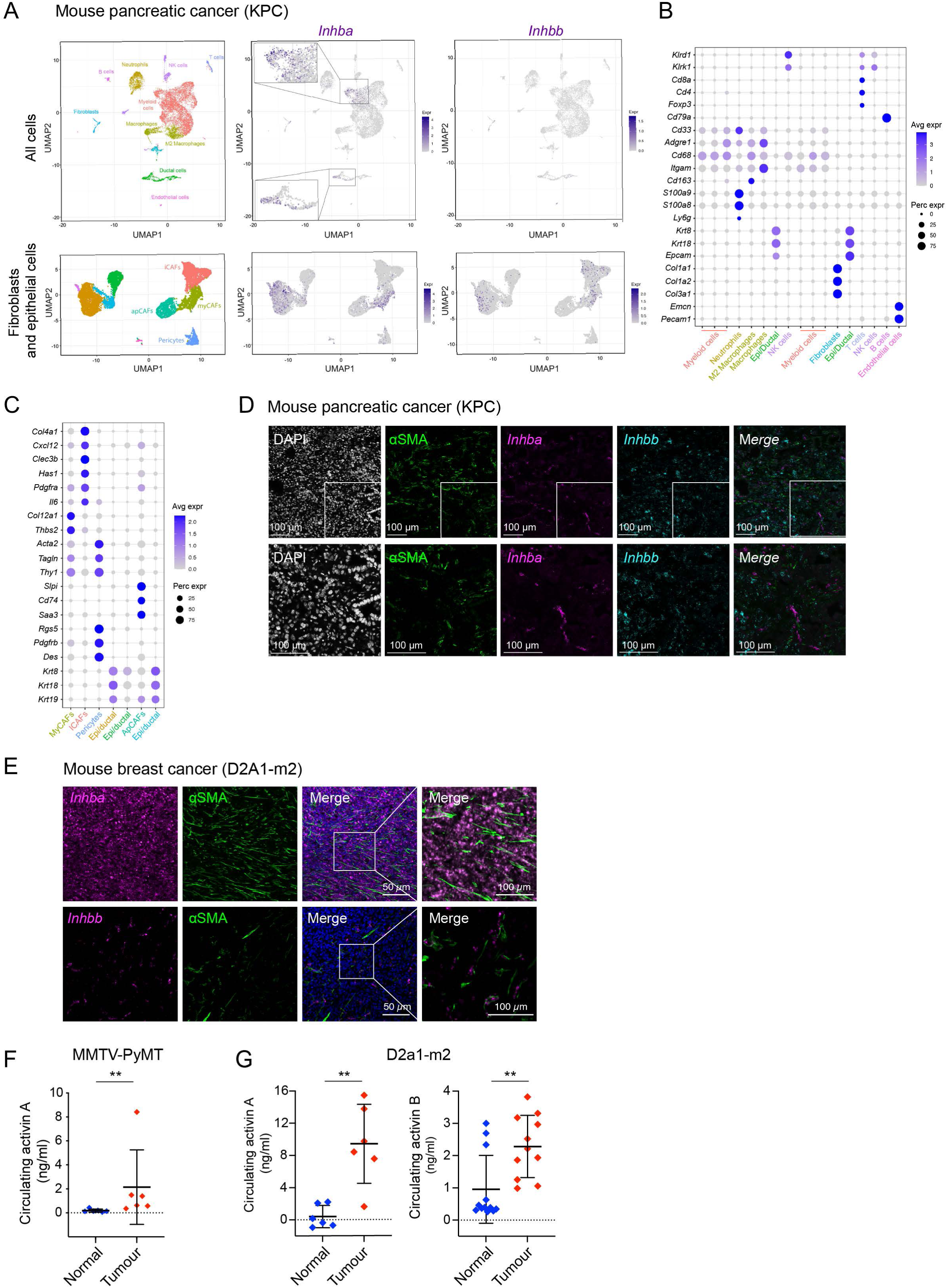
Activin A and activin B are expressed in mouse models of pancreatic and breast tumours. (**A**) UMAP projections from a re-analyzed mouse scRNA-seq dataset ^125^ displaying the viable cell population (upper panels) or fibroblasts and epithelial cells (lower panels), with all cell types and indicating levels of *Inhba* and *Inhbb* expression. The *Inhba* panel is further illustrated with zoomed insets to highlight the highly expressing cells. (**B, C**) Dot plots demonstrating representative marker genes across all cell types (B) and specifically fibroblasts and epithelial cells (C). Dot size is proportional to the fraction of cells expressing specific genes. Colour intensity corresponds to the relative expression of specific genes. (**D**) Confocal microscopy images of mouse PDAC samples (KPC) stained for *Inhba* (magenta) and *Inhbb* (cyan) mRNA by RNAScope™ and for αSMA protein (green) by immunofluorescence with nuclei visualized by DAPI staining (white). The bottom row shows higher magnifications of the regions marked with a white box. Scale bars correspond to 100 µm. (**E**) Confocal microscopy images of mouse breast cancer samples (D2A1-m2 model) stained for *Inhba* mRNA (magenta) (upper panels) or *Inhbb* mRNA (magenta) by RNAScope (lower panels) and for αSMA protein (green) by immunofluorescence with nuclei visualized by DAPI staining (blue). A higher magnification is shown for the merge panels. The scale bars correspond to 50 µm and 100 µm as indicated. (**F**) Graph demonstrating circulating activin A levels in the MMTV-PyMT breast cancer mouse model comparing tumour-bearing mice and healthy controls. Means ± SD are shown. **, p <0.01. (**G**) Graphs demonstrating circulating activin A (left graph) and activin B (right graph) levels in the D2A1-m2 breast cancer mouse model, comparing tumour bearing mice and healthy controls. Means ± SD are shown. **, p <0.01. Note that some of the values for activin A in the normal mice were below the detection of the plate reader.

**Figure S2.**
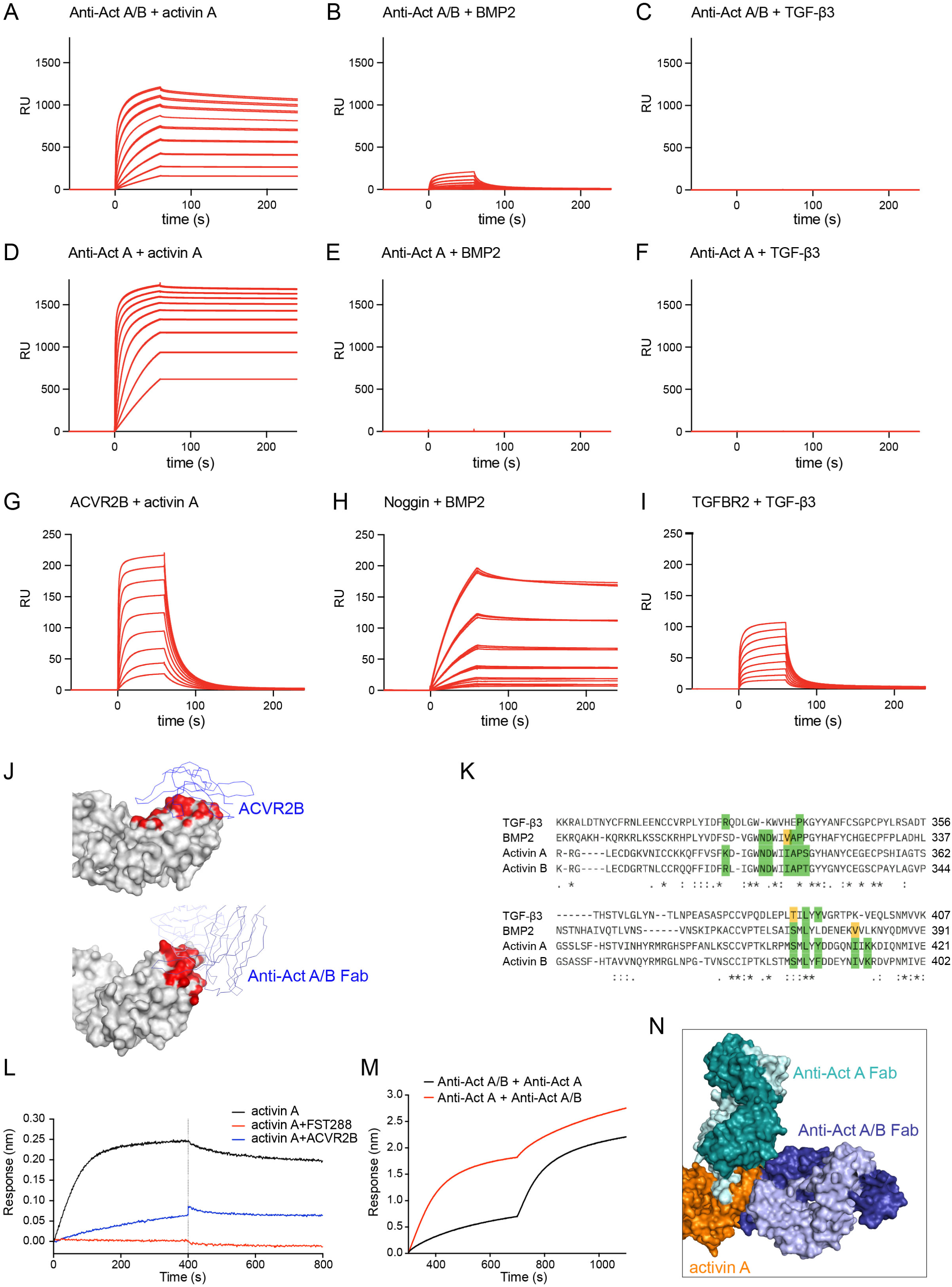
Characterization of the anti-activin A/B antibody. (**A–C**) SPR binding sensorgrams of Anti-Act A/B binding to immobilized activin A (A), BMP2 (B), or TGF-β3 (C) on the chip surface. The concentrations of anti-Act A/B ranged from 2 µM to 7.8 nM. The K_D_ for Anti-Act A/B binding activin A was calculated to be 0.254 nM; K_D_ for anti-Act A/B binding BMP2 was 0.102 µM and no binding was detected for anti-Act A/B binding TGF-β3. (**D–F**) SPR binding sensorgrams of anti-Act A binding to immobilized activin A (A), BMP2 (B), or TGF-β3 (C) on the chip surface. The concentrations of anti-Act A ranged from 2 µM to 7.8 nM. The K_D_ for anti-Act A binding activin A was calculated to be 8.74 pM. No binding was detected for anti-Act A binding BMP2 or TGF-β3. (**G**) Controls for panels A and D. SPR binding sensorgrams of ACVR2B extracellular domain (ECD) binding to immobilized activin A. The concentrations of ACVR2B ECD ranged from 2 µM to 7.8 nM. The K_D_ was calculated as 0.114 µM. (**H**) Controls for panels B and E. SPR binding sensorgrams of noggin binding to immobilized BMP2. The concentrations of Noggin ranged from 62.5 nM to 2 nM. The K_D_ was calculated as 1.26 nM. (**I**) Controls for panels C and F. SPR binding sensorgrams of TGFBR2 extracellular domain (ECD) binding to immobilized TGF-β3. The concentrations of TGFBR2 ECD ranged from 4 µM to 15.63 nM. The K_D_ was calculated as 0.317 µM. In all cases, for each concentration of ligand binder (antibody or control) duplicates are shown. (**J**) Comparison of structure of activin A with the extracellular domain of ACVR2B (PDB:1s4y) and AlphaFold 3 model of activin A and anti-activin A/B Fab. Activin A proteins are shown with molecular surface where atoms within 4.0 Å from the ACVR2B or the anti-activin A/B Fab are coloured red, showing significant overlap between the two. The ACVR2B or the anti-activin A/B Fab structures are shown as Cα traces in blue. (**K**) Alignment of mature TGF-β3, BMP2 activin A and activin B to show how the interacting residues are conserved between activin A and activin B, but less so between activin A/B and TGF-β3 and BMP2, explaining the specificity of antibody binding. Residues coloured green are anti-activin A/B interacting residues that are identical to those in activin A and residues coloured yellow are similar in chemical properties. (**L**) The anti-activin A/B antibody competes with follistatin (FST288) and the extracellular domain of ACVR2B for binding to activin A as measured by bio-layer interferometry. (**M**) Bio-layer interferometry reveals that the anti-activin A/B and anti-activin A Fab fragments do not compete for binding to activin A. The same result is observed regardless of the order of addition of the different Fab fragments. (**N**) Comparison of anti-activin A/B and anti-activin A Fab fragments in complex with activin A using AlphaFold3. Activin A (orange) models in complex with the anti-activin A/B Fab fragment (dark and light blue) and anti-activin A antibody Fab fragment (dark and light teal) showing non-overlapping binding sites.

**Figure S3.**
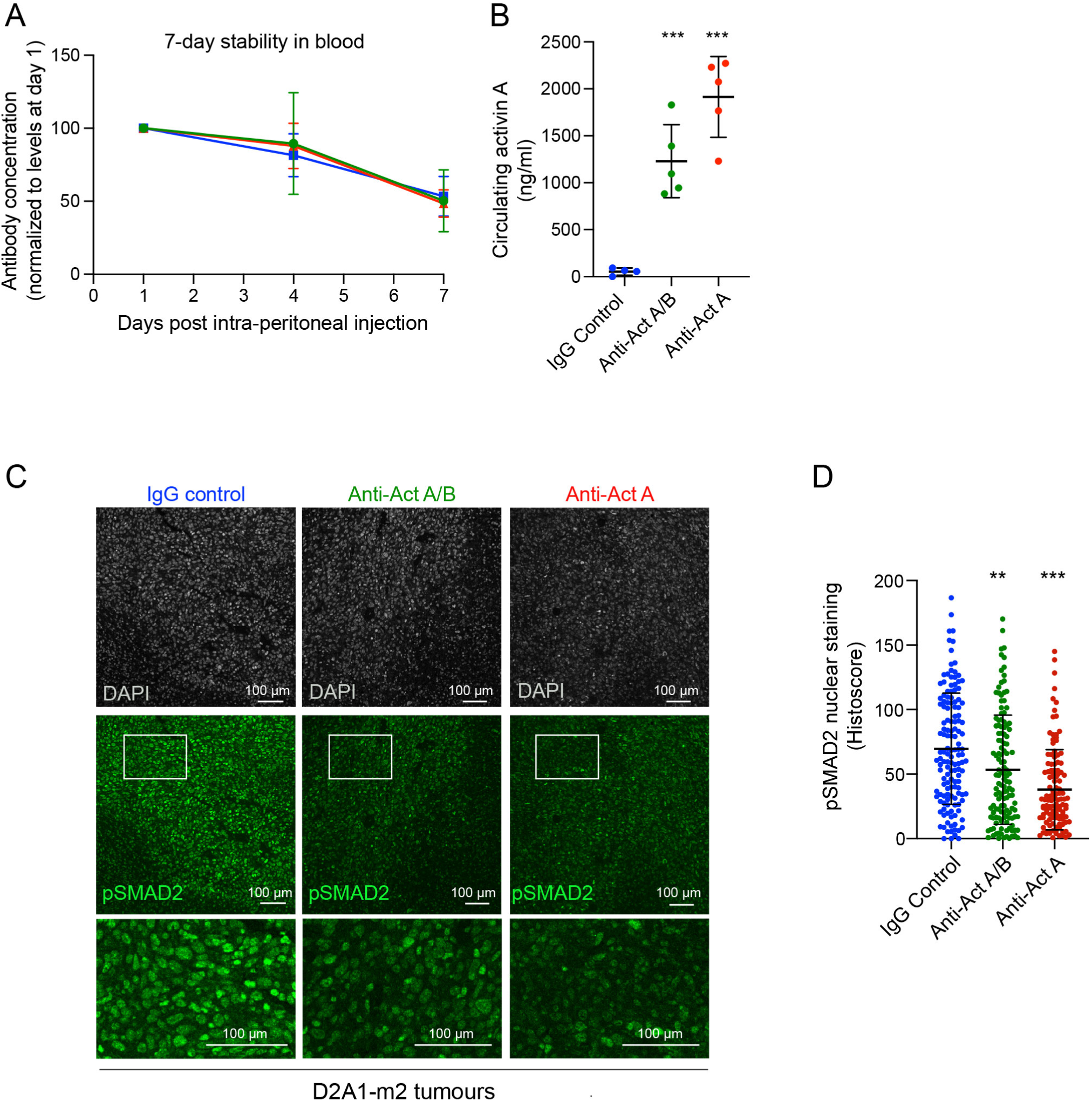
The anti-activin antibodies are stable in the bloodstream and effectively neutralize activin activity. (**A**) Graph demonstrating the stability of the IgG control, anti-activin A/B and anti-activin A antibodies in the blood of injected FvB/NJ mice, measured by ELISA against human IgG. The levels of each antibody are normalized to 100 at day 1 post intra-peritoneal injection. Means ± SD are shown. (**B**) Circulating levels of activin A, measured by ELISA from mice treated with the IgG control, anti-activin A/B and anti-activin A antibodies. Means and SD are shown. ***, p <0.001. (**C**) Representative confocal microscopy images of pSMAD2 staining in mouse breast tumours from the D2A1-m2 model. Mice either treated with either anti-activin A, anti-activin A/B or IgG control antibodies. Nuclei are stained with Dapi (grey). A higher magnification of the area in the white boxes is shown below. Scale bar corresponds to 100 µm. (**D**) Quantification of pSMAD2 staining from the experiment shown in panel C using the Histoscore method. The Histoscore was calculated according to the following formula: Histoscore = [(0 x % negative cells) + (1 x % weak positive cells) + (2 x % moderate positive cells) + (3 x % strong positive cells), with the overall score ranging from 0 (negative) to 300 (100% strong staining)]. Three tumours per condition were analysed with 40 fields of view per tumour. Means ± SD are shown. **, p <0.01; ***, P <0.001.

**Figure S4.**
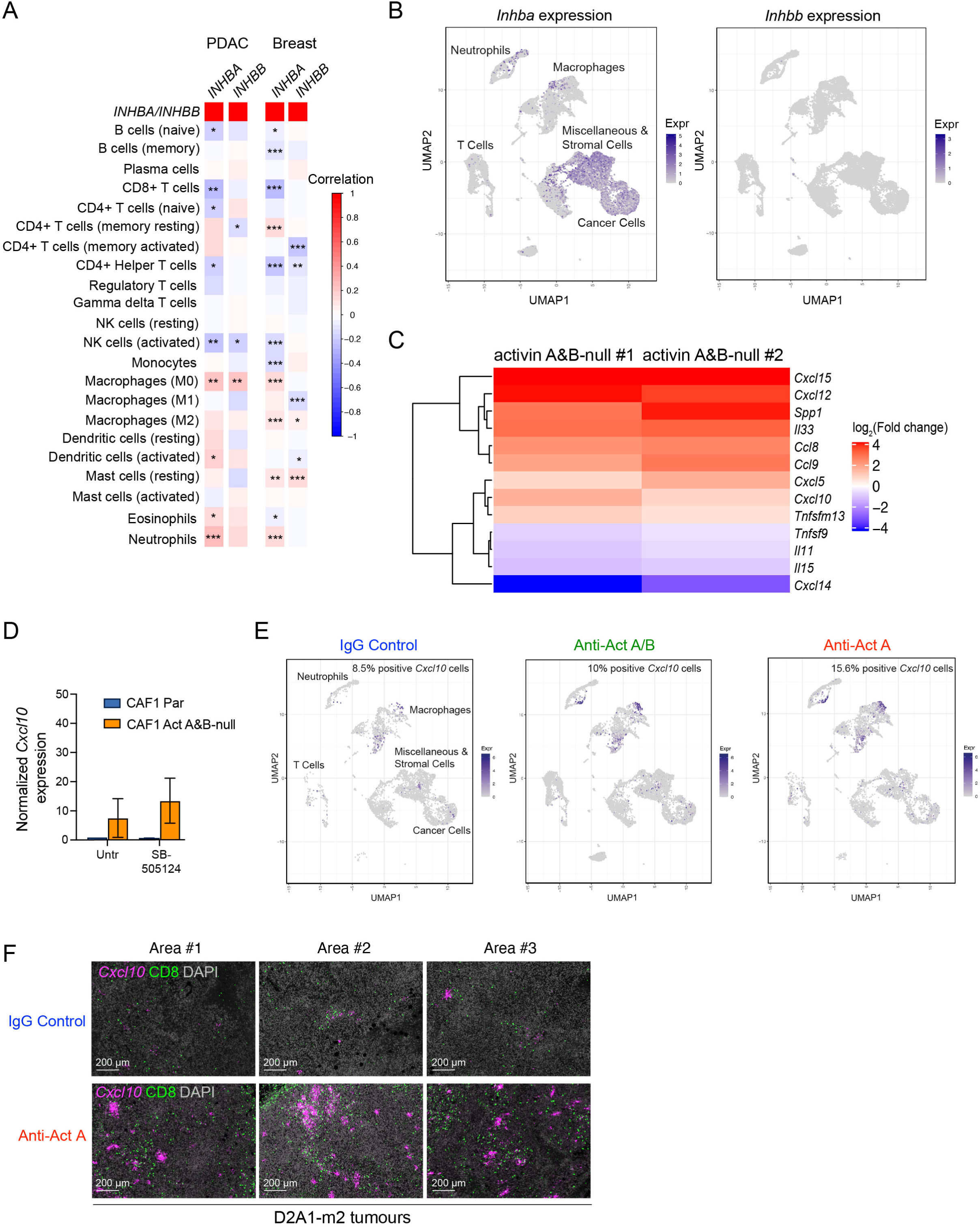
Expression of *INHBA* negatively correlates with the presence of CD8+ T cells. (**A**) CIBERSORT immune deconvolution of human pancreatic (left) and breast (right) TCGA cohorts showing correlations between *INHBA* expression with different immune cell types. Statistical significance is indicated. *, p< 0.05, **, p <0.005; ***, P <0.0005. (**B**) UMAP projections from an scRNA-seq dataset of tumours from the D2A1-m2 mouse breast cancer model (D2A1-m2 injected into the mammary fat pad of BALB/c mice), as in Figure 4A, indicating *Inhba* and *Inhbb* expression. All treatment groups were collated for each UMAP. (**C**) A heatmap of a manually curated list of 13 genes associated with immune cell modulation that shows significant differential expression between two activin A&B-null CAF1 clones and parental CAF1 cells. The data are represented as the log_2_Fc expression of each gene in each Activin A/B-null CAF1 clone compared to the parental CAF1 cells. (**D**) Expression levels of *Cxcl10* measured by qPCR in parental (Par) CAF1 cells and ActivinA&B-null CAF1 cells. Cells either untreated (Untr) or treated with TGF-β/activin receptor inhibitor SB-505124. Relative gene expression was calculated by normalizing to the Untr group and to *Gapdh* mRNA levels for each sample. The data are represented as the mean ± SD of 2 biological replicates. (**E**) UMAP projections from an scRNA-seq dataset of tumours from the D2A1-m2 mouse breast cancer model (D2A1-m2 injected into the mammary fat pad of BALB/c mice), as in Figure 4A indicating *Cxcl10* expression for the mice treated with IgG control, anti-activin A/B and anti-activin A antibodies. (**F**) Three representative examples of *Cxcl10* RNAscope™ staining multiplexed with CD8 staining on D2A1-m2 tumours from mice treated with either IgG control or the anti-activin A antibody. Scale bar, 200 µm.

**Figure S5.**
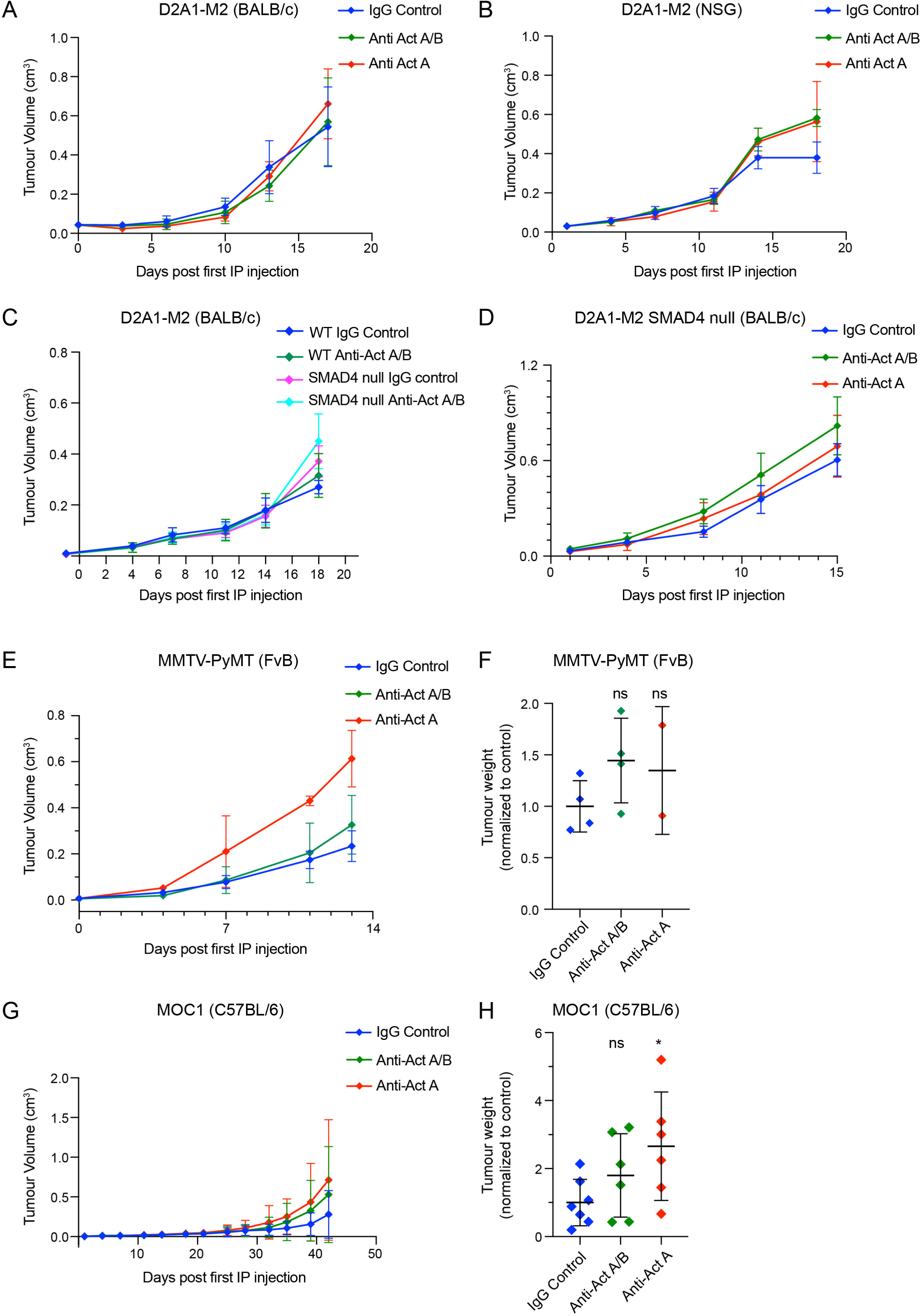
Treatment of mice with the anti-activin antibodies leads to a small increase in tumour size as measured by tumour volume. (**A–D**) Tumour volumes over time relative to the day of the first antibody injection for the experiments shown in Figure 5 (A–D). D2A1-m2 cells (wild type or SMAD4-null as indicated) were injected into the mammary fat pad of BALB/c or NSG mice as indicated. Mice were treated with IgG control, anti-activin A/B or anti-activin A antibodies and the tumour volumes were measured over time. (**E, F**) Tumour volumes over time (E) and final tumour weights (F) in the MMTV-PyMT mouse breast cancer model (tumour cells isolated from a MMTV-PyMT mouse injected into the mammary fat pad of FvB/NJ mice). Mice were treated with IgG control, anti-activin A/B or anti-activin A antibodies. ns, not significant (**G, H**) Tumour volumes over time (G) and final tumour weights (H) in the MOC1 mouse oral squamous cell carcinoma (MOC1 cells injected into subcutaneously into C57BL/6J mice). Mice were treated with IgG control, anti-activin A/B or anti-activin A antibodies. ns, not significant; *, p < 0.05. The statistical significance is calculated relative to the IgG control.

**Figure S6.**
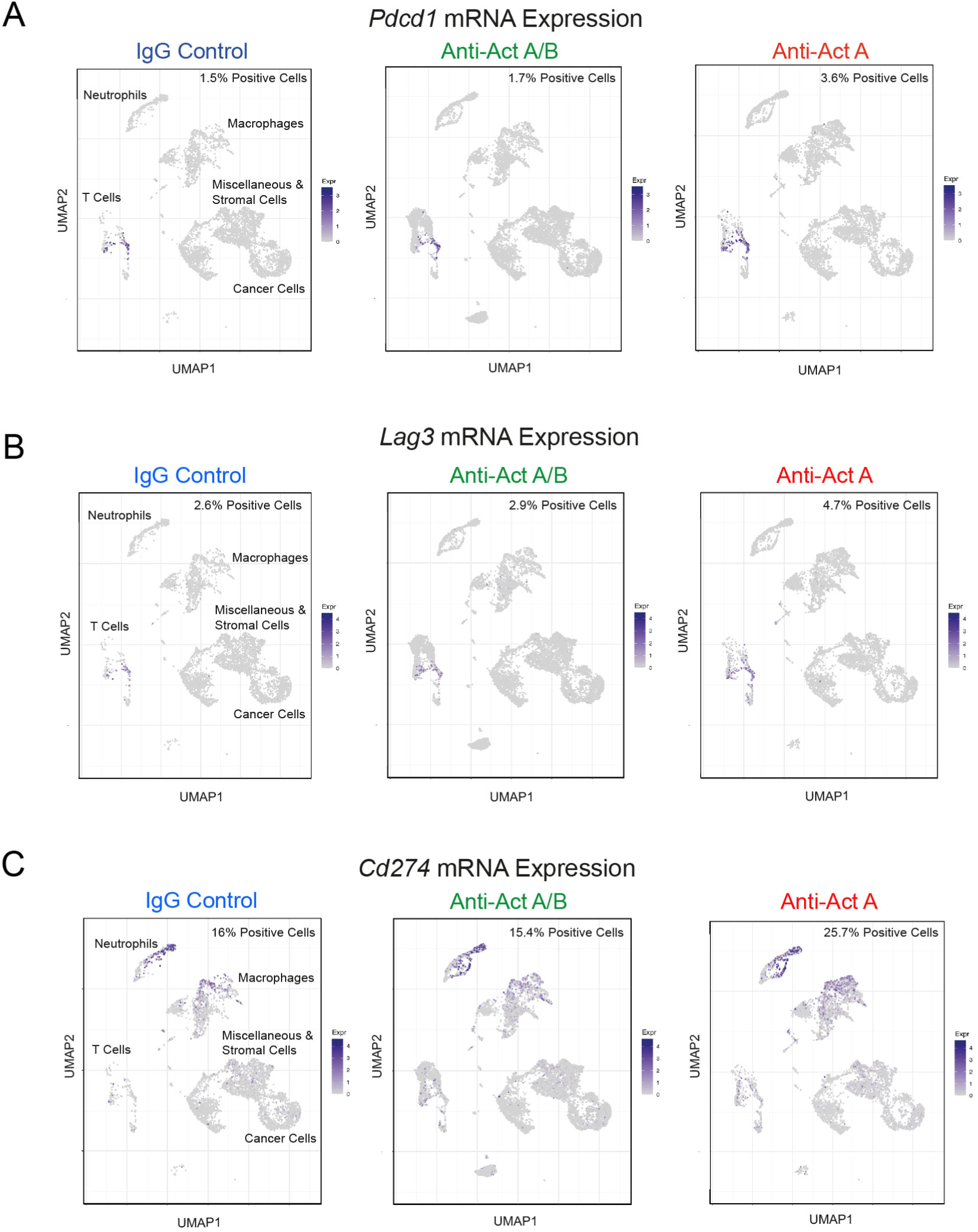
Treatment with the anti-activin A antibody is associated with upregulation of exhaustion markers PD-1 and LAG3 in infiltrating T cells. (**A–C**) UMAP projections from the scRNA-seq dataset of tumours from the D2A1-m2 mouse breast cancer model (D2A1-m2 injected into the mammary fat pad of BALB/c mice), as in Figure 4A indicating *Pdcd1* expression (gene encoding PD1) (A), *Lag3* expression (B) and *Cd274* expression (gene encoding PD-L1) (C), for the mice treated with IgG control, anti-activin A/B and anti-activin A antibodies.

**Figure S7.**
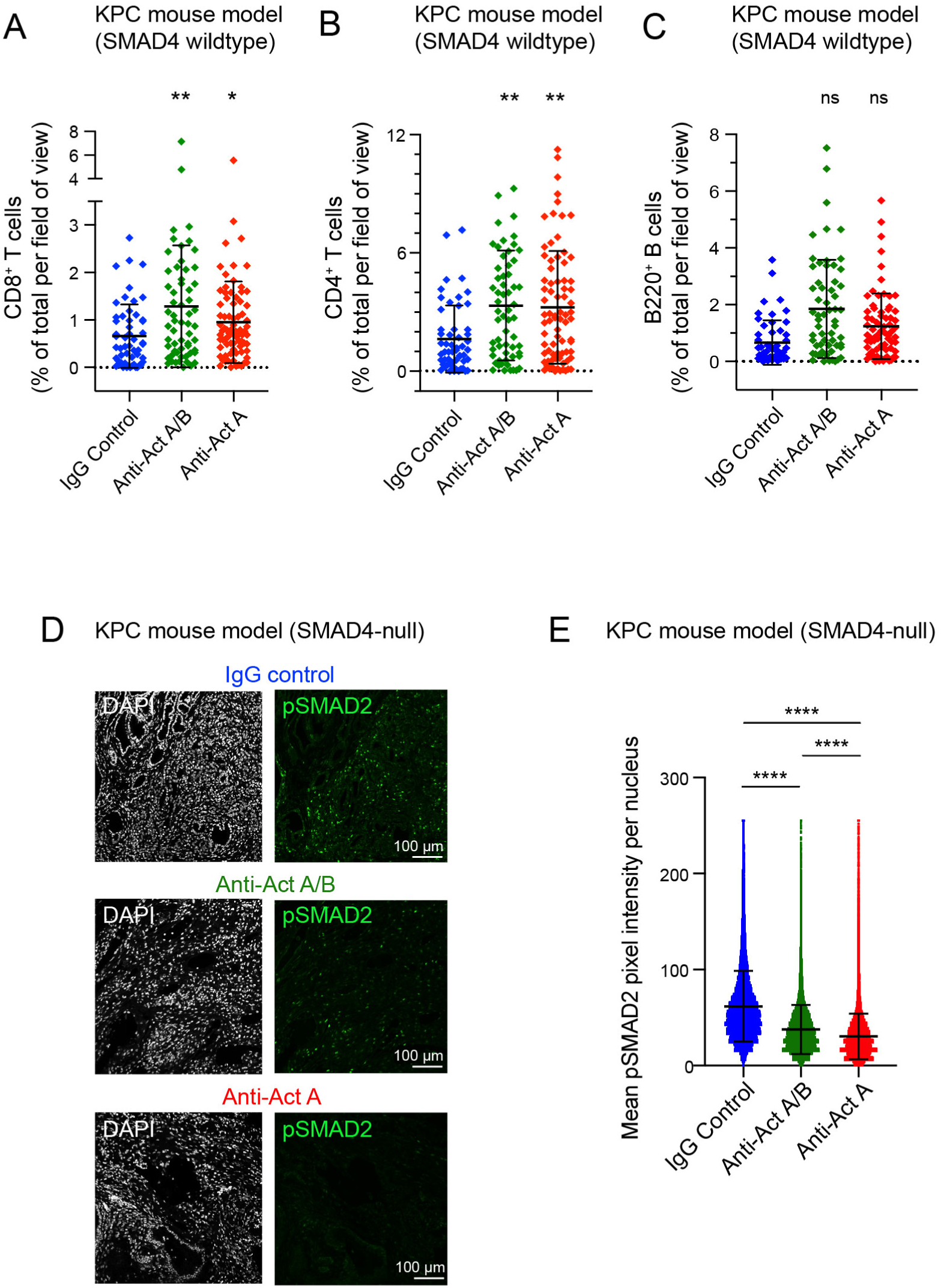
Treatment of KPC PDAC mice with anti-activin antibodies leads to an increase in cytotoxic T cell infiltration. (**A–C**) Quantifications of immune cell infiltration in the mouse pancreatic cancer model (KPC) treated with IgG control anti-activin A/B or anti-activin A antibodies. CD8^+^ (A), CD4^+^ (B) and B220^+^ (C) cells were analysed. The following numbers of tumours were analysed: IgG control (n=5), anti-activin A/B (n=6), anti-activin A (n=7). The data are represented as % of total positive cells, per field of view. 10 fields of view were analysed per tumour. ns, not significant; *, p, 0.05; **, p <0.01. (**D**) Confocal microscopy images of pSMAD2 staining in tumours from the SMAD4-null PDAC mouse model model. Mice either treated with either anti-activin A, anti-activin A/B or IgG control antibodies. DAPI (grey)and pSMAD2 (green) channels shown for each treatment group. Scale bar corresponds to 100 μm. (**E**) Quantification of pSMAD2 staining in tumours from the SMAD4-null PDAC mouse model model. Mice either treated with either anti-activin A, anti-activin A/B or IgG control antibodies. Data represented are the mean pixel intensity of pSMAD2 staining per nuclei. 2 tumours per condition with 6 FOV per tumour. Means ± SD are shown. ****, p <0.0001.

**Figure S8.**
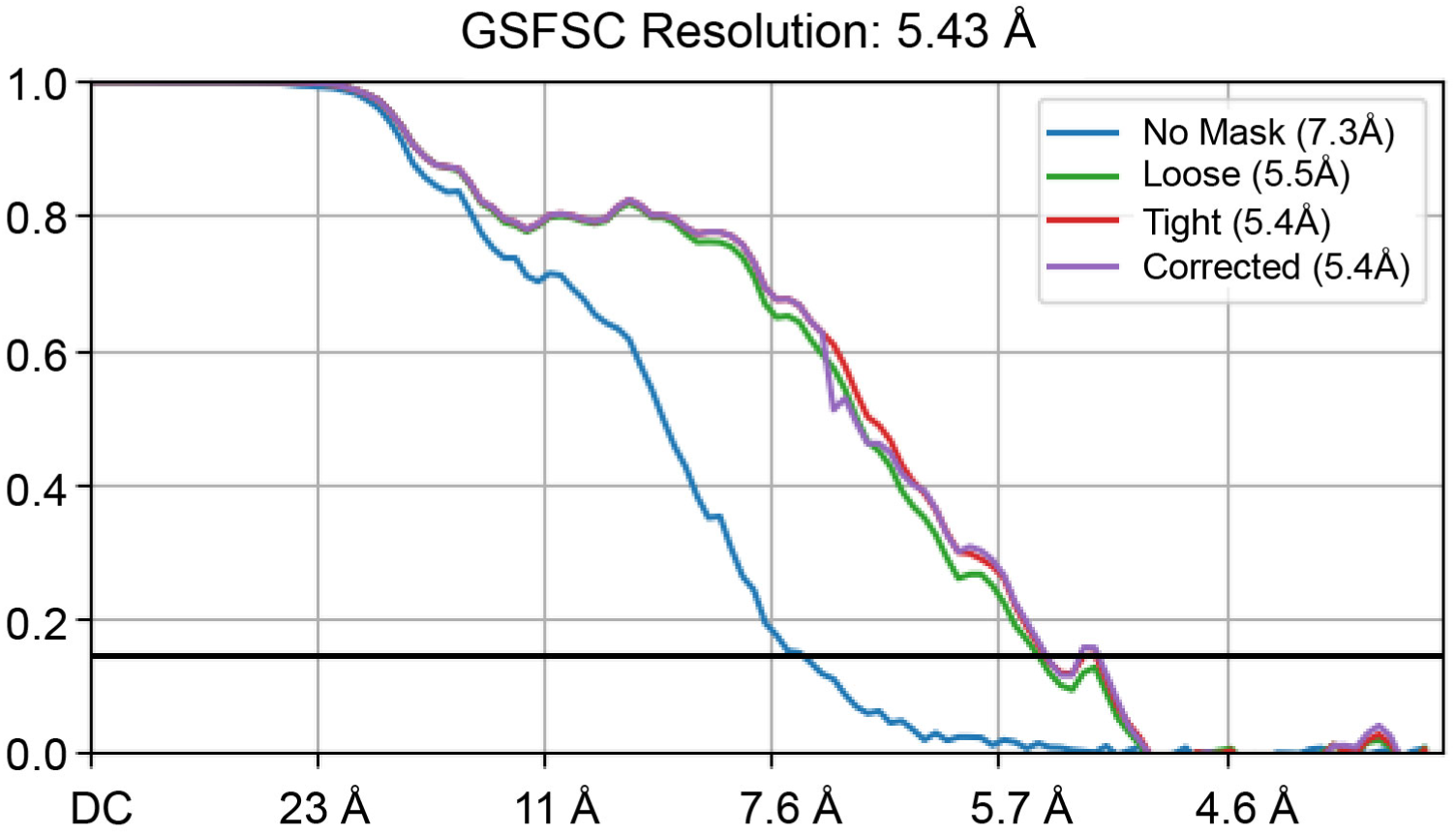
Global GSFSC curve for the C1 map for activin A–anti-activin A/B Fab complex.

**Supplementary Table 1.**
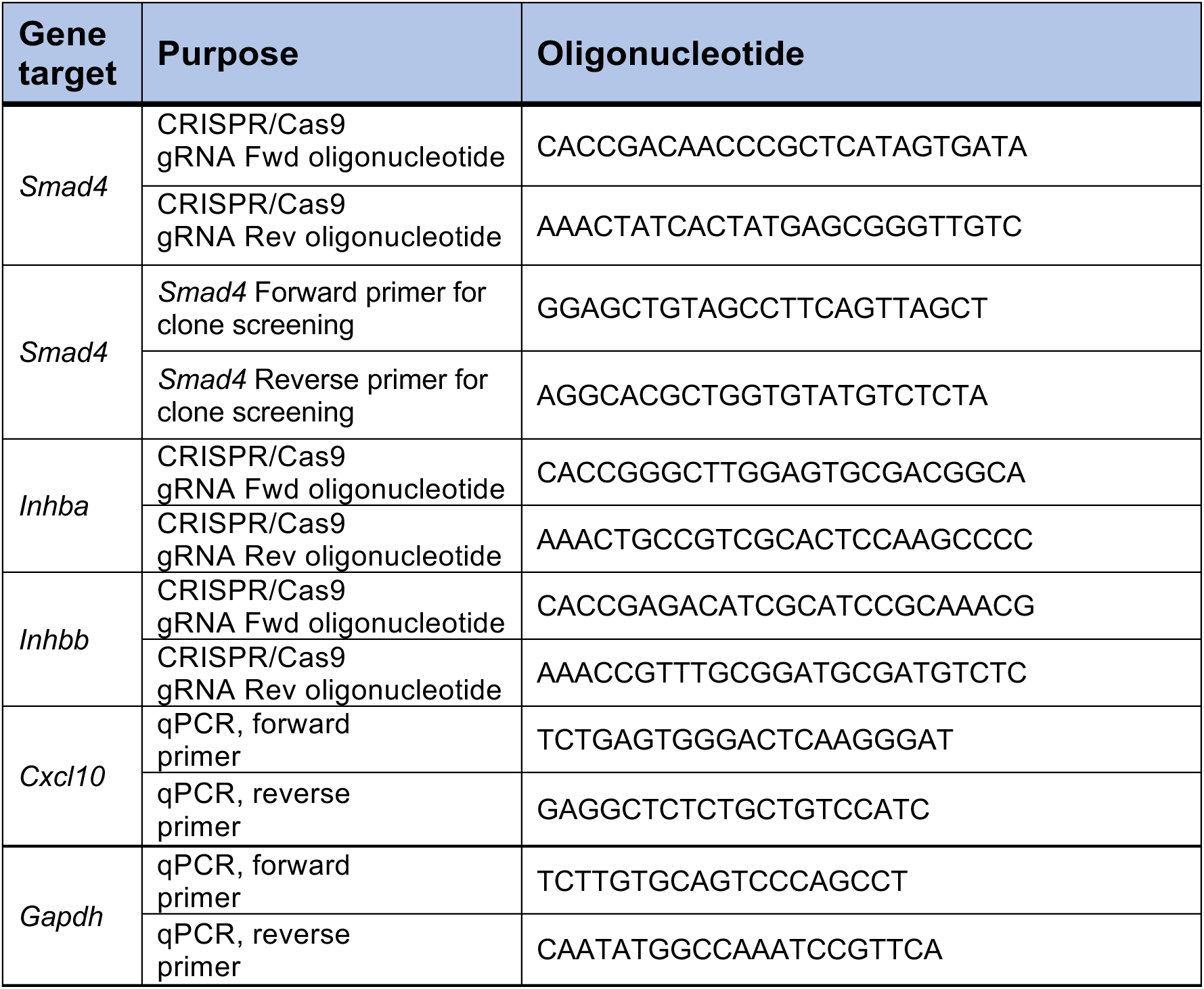
Oligonucleotides.

**Supplementary Table 2.** Key Resources.

| REAGENT or RESOURCE | SOURCE | IDENTIFIER |
| --- | --- | --- |
| **Antibodies** |  |  |
| Anti-phospho-Smad2 (Western and IF) | Cell Signaling Technology | Cat # 3108<br>RRID:AB_490941 |
| Anti-activin A (Western) | Santa Cruz Biotechnology | Cat # sc-166503<br>RRID:AB_2125854 |
| Anti-activin B (Western) | Santa Cruz Biotechnology | Cat # sc-376971<br>RRID:AB_3677370 |
| Anti-SMAD2/3 (Western) | BD Biosciences | Cat # 610842<br>RRID:AB_398161 |
| Anti-Tubulin (Western) | Abcam | Cat # Ab6160;<br>RRID:AB_305328 |
| Anti-P-MYL9 (Western) | Cell Signaling Technology | Cat # 3674<br>RRID:AB_2147464 |
| Anti-MLC (Western) | Cell Signaling Technology | Cat # 3672<br>RRID:AB_10692513 |
| Anti-αSMA (IF) | Agilent | Cat # M0851<br>RRID:AB_2223500 |
| Anti-CD4 (IHC) | Abcam | Cat # ab183685<br>RRID:AB_2686917 |
| Anti-CD8 (IHC) | Ebiosciences | Cat # 14-0808-82<br>RRID:AB_2572861 |
| Biotin anti-mouse CD45R/B220 (IHC) | BD Biosciences | Cat # 553086<br>RRID:AB_394616 |
| Anti-Ki67 (IHC) | Abcam | Cat # ab15580<br>RRID:AB_443209 |
| Anti-active caspase 3 (IHC) | R&D Systems | Cat # AF835<br>RRID:AB_2243952 |
| HRP-conjugated anti-mouse secondary antibody | Dako | Cat # P0447<br>RRID: AB_2617137 |
| HRP-conjugated anti-rabbit secondary antibody | Dako | Cat # P0448<br>RRID: AB_2617138 |
| HRP-conjugated anti-rat secondary antibody | Jackson ImmunoResearch | Cat # 712-035-153<br>RRID:AB_2340639 |
| Alexa Fluor-594-conjugated donkey anti-mouse secondary antibody. | Invitrogen | Cat # A21203<br>RRID:AB_2535789 |
| Biotinylated donkey-anti rabbit secondary antibody | Jackson ImmunoResearch | Cat # 711-065-152<br>RRID:AB_2340593 |
| Biotinylated goat anti-rabbit secondary antibody | Vector Laboratories | Cat # BA-1000<br>RRID:AB_2313606 |
| Biotinylated goat anti-rat secondary antibody | Vector Laboratories | Cat # BA-4001<br>RRID:AB_10015300 |
| Anti-M13 HRP-conjugated antibody | Sino Biological | Cat # 11973-MM05T-H<br>RRID:AB_2857928 |
| **Chemicals, peptides, and recombinant proteins** |  |  |
| Streptavidin Alexa Fluor-488 | Molecular Probes | Cat # S32354 |
| SB-505124 | Tocris | Cat # 3263 |
| PBS | ThermoFisher Scientific | Cat # 10091403 |
| DMEM | Gibco | Cat # 11995065 |
| Collagenase D | Roche | Cat # 11088866001 |
| Dispase II | Roche | Cat # 4942078001 |
| DNAase I | Sigma-Aldrich | Cat # D5025-375K |
| Hanks Balanced Salt Solution (HBSS) | Gibco | Cat # 14025092 |
| BSA | Sigma-Aldrich | Cat # A9418 |
| RBC Lysis Buffer | Biolegend | Cat # 420301 |
| Growth factor reduced (GFR) Matrigel | Corning | Cat # 356231 |
| IPTG | Sigma | Cat # I6758 |
| DAPI | Sigma | Cat # 10236276001 |
| FuGene® 6 | Promega | Cat # E269A |
| Fetal Bovine Serum | ThermoFisher Scientific | Cat # 10270-106 |
| TRIzol | Invitrogen | Cat # 15596026 |
| Activin A for cell inductions | PeproTech | Cat #120-14 |
| TGF-β1 for cell inductions | PeproTech | Cat #100-21C |
| BMP2 | PeproTech | Cat # 120-02 |
| Noggin | PeproTech | Cat # 120-10C |
| Follistatin | Sigma | Cat # F2177 |
| TGF-β3 | Ref<sup>1</sup> | N/A |
| TGF-β blocking antibody (LY2424087) | Eli Lilly | N/A |
| ACVR2B-ECD | This paper | N/A |
| ACVR2B-ECD | Ref<sup>1</sup> | N/A |
| TGFBR2-ECD | Ref<sup>2</sup> | N/A |
| Human anti-activin A/B antibody | This paper | N/A |
| Human anti-activin A antibody | This paper | N/A |
| Human anti-NIP228 IgG control antibody | Ref<sup>3</sup> | N/A |
| Mouse anti-PD-L1 D265A clone 80 | Ref<sup>4</sup> | N/A |
| Rat anti-PD-L1 clone 10F.9G2 | Ref<sup>4</sup> | N/A |
| Mouse anti-NIP228 D265 IgG control | Ref<sup>4</sup> | N/A |
| Rat anti-NIP228 IgG2b K control | Ref<sup>4</sup> | N/A |
| **Critical Commercial Assays** |  |  |
| RNAscope Multiplex Fluorescent Reagent kit v2 | ACDBio | Cat # 323100) |
| polyA KAPA mRNA HyperPrep kit | KAPA Biosystems | Cat # KK8580 |
| TSA Plus Biotin Systems | Akoya | Cat #<br>NEL749A001KT |
| ELISA assay for mouse activin A | Ansh Lab | Cat # AL-150 |
| ELISA assay for mouse activin B | Ansh Lab | Cat # AL-156 |
| ELISA assay for human/mouse activin A | R&D Systems | Cat # DAC00B |
| ELISA for human IgG1 | ThermoFisher Scientific | Cat # EHIGG1 |
| VECTASTAIN ABC-HRP kit | Vector Laboratories | Cat # PK-4000 |
| EZ-Link™ NHS-PEG4-Biotin | Thermo Scientific | Cat # A39259 |
| Picrosirius Red kit | Abcam | Cat # ab150681 |
| RNeasy kit | Qiagen | Cat # 74104 |
| **Experimental models: Cell Lines** |  |  |
| D2A1-m2 | Clare Isacke (ICR)<sup>5</sup> | N/A |
| D2A1-m2 SMAD4-null | This paper | N/A |
| HaCaT | Francis Crick Institute (FCI) Cell Services | N/A |
| HEK293T CAGA<sub>12</sub>-Luciferase/Renilla | FCI Cell Services<sup>6</sup> | N/A |
| CAF1 | Erik Sahai (FCI)<sup>7</sup> | N/A |
| CAF1 Activin A&B Null clone 1 | This paper | N/A |
| CAF1 Activin A&B Null clone 2 | This paper | N/A |
| NF1 | Erik Sahai (FCI)<sup>7</sup> | N/A |
| MOC1 | Erik Sahai (FCI)<sup>8</sup> | N/A |
| FreeStyle™ 293-F | Francis Crick Institute (FCI) Cell Services | N/A |
| Primary MMTV-PyMT cancer cells | Prepared from tumours of MMTV-PyMT mice. | N/A |
| **Experimental models: Organisms/Strains** |  |  |
| *Mus Musculus* BALB/cJ | JAX strain #000651 | https://www.jax.org/strain/000651 |
| *Mus Musculus* FVB/NJ | JAX strain #001800 | https://www.jax.org/strain/001800 |
| *Mus Musculus* C57BL/6J | JAX strain #000664 | https://www.jax.org/strain/000664 |
| *Mus Musculus* NOD.Cg-Prkdc<sup>scid</sup> Il2rg<sup>tm1Wjl</sup>/SzJ | Ref <sup>9</sup> |  |
| *Mus Musculus* Trp53<sup>loxP/LoxP</sup> | Ref <sup>10</sup> |  |
| *Mus Musculus* Tg(Pdx1-cre)6Tuv | Ref <sup>11</sup> |  |
| *Mus Musculus* Ptf1a<sup>tm1.1(cre)Cvw</sup> | Ref <sup>12</sup> |  |
| *Mus Musculus* Smad4<sup>tm1Rob/tm1Rob</sup> | Ref <sup>13</sup> |  |
| *Mus Musculus* Kras<sup>LSL-G12D/+</sup> | Ref <sup>14</sup> |  |
| *Mus Musculus* Trp53<sup>LSL-R172H/+</sup> | Ref <sup>15</sup> |  |
| **Recombinant DNA** |  |  |
| PX458 | Ref <sup>16</sup> | Addgene plasmid #48138 |
| pVITRO1-dV-IgG1/lambda | Ref <sup>17</sup> | Addgene plasmid #52214 |
| pOP3BT | Marko Hyvonen Lab | Addgene #112603 |
| pBAT4 | Marko Hyvonen Lab | Addgene #112580 |
| **Software and algorithms** |  |  |
| FIJI (ImageJ) | Ref <sup>18</sup> | https://imagej.net/Fiji/Downloads |
| QuPath | QuPath | https://qupath.github.io/ |
| GraphPad Prism 10 | GraphPad | https://www.graphpad.com/scientific-software/prism/ |
| AlphaFold3 | Ref <sup>19</sup> | (https://alphafoldserver.com) |
| CryoSPARC V4 | Ref <sup>20</sup> | https://cryosparc.com |
| cutadapt-1.9.1 | Ref <sup>21</sup> |  |
| RSEM-1.3.0/STAR-2.5.2 | Ref <sup>22,23</sup> |  |
| DESeq2 (version 1.24.0) | Ref <sup>24</sup> |  |
| ClusterProfiler package (version 3.12.0) | Ref <sup>25</sup> |  |
| ReactomePA package (version 1.28.0) | Ref <sup>26</sup> |  |
| ComplexHeatmap package (version 2.0.0) | Ref <sup>27</sup> |  |
| Seurat R-package (version 4.0.5) | Ref <sup>28</sup> |  |
| glmGamPoi R-package, version 1.1.13 | Ref <sup>29</sup> |  |
| **Other** |  |  |
| (RNAscope) mouse *Inhba* | ACDbio | Cat # 455871-C2 |
| (RNAscope) mouse *Inhbb* | ACDbio | Cat # 475271-C1 |
| (RNAscope) human *INHBA* | ACDbio | Cat # 415111-C2 |
| (RNAscope) human *INHBB* | ACDbio | Cat # 435741-C1 |
| RNAscope mouse *Cxcl10* | ACDbio | Cat # 408921-C2 |

